# Modeling the inverse MEG problem in neuro-imaging using Physics Informed Neural Networks

**DOI:** 10.64898/2026.02.03.703484

**Authors:** Ourania Giannopoulou

## Abstract

Magnetoencephalography (MEG) forward and inverse modeling is fundamental to neuroscientific discovery, yet the inversion of partial differential equations (PDEs) remains one of the most difficult challenges due to its inherent ill-posedness. While traditional numerical methods often struggle with the computational burden and regularization requirements of these problems, neural networks have recently emerged as a highly viable alternative, offering the ability to learn complex, non-linear mappings and provide efficient, real-time inference. This paper presents a framework for the MEG forward and inverse problems, integrating finite element modeling with neural network techniques. The forward problem is solved using FEniCS to model the electric potential governed by the Poisson equation on a realistic anatomical brain mesh, with magnetic fields computed via the Biot-Savart law. For the inverse problem, we introduce a Physics-Informed Neural Network (PINN) approach in order to deal with the ill condition of the problem. Unlike purely data-driven deep learning approaches that treat this problem as a black box learned from massive datasets, the proposed PINN framework directly embeds the governing physics—Maxwell’s equations and the Biot-Savart law—into the loss function, ensuring that the reconstructed sources satisfy the fundamental electromagnetic laws even in data-scarce regimes. We validate the framework on a high-resolution anatomical mesh and compared against the standard Minimum Norm Estimation (MNE). Results demonstrate that the PINN approach achieves a 30.2% improvement over the MNE baseline.

## 1 Introduction

### 1.1 Magnetoencephalography and its Applications

Magnetoencephalography (MEG) is a powerful, non-invasive neuroimaging technique that measures the magnetic fields produced by electrical currents in neurons (Hämäläinen et al. 1993; Ebersole and Ebersole 2010). Unlike electroencephalography (EEG), which measures electric potentials on the scalp (Teplan 2002), MEG detects magnetic fields that are less distorted by the skull’s low conductivity, offering superior spatial resolution for localizing neural activity (Hämäläinen et al. 1993). A fundamental advantage of MEG lies in its sensitivity to source orientation; it primarily detects the tangential components of primary currents, making it complementary to EEG which is more sensitive to radial sources (Sarvas 1987).

MEG has found widespread application in both clinical and research settings (Baillet et al. 2001). Clinically, it is indispensable for pre-surgical epilepsy mapping, where it localizes epileptogenic foci, and for functional mapping of eloquent cortex (motor, sensory, language areas) prior to tumor resection. In research, MEG enables the study of cognitive processes with millisecond temporal resolution, elucidating neural oscillations, functional connectivity, and event-related responses (Picton et al. 2000). However, realizing the full potential of MEG requires solving the challenging inverse problem: reconstructing the 3D internal current sources from 2D sensor measurements.

### 1.2 Historical Development and Geometric Modeling

The mathematical foundations of MEG were established in the 1960s and 1970s by Geselowitz (Geselowitz 1967, 1970), who derived the fundamental relationships between internal current sources and external fields in volume conductors. Seminally, Sarvas (1987) provided an exact analytical solution for the magnetic field of a current dipole in a spherical conductor, which remains a standard benchmark for forward modeling.

While spherical models offer computational simplicity, they oversimplify the complex non-spherical geometry of the human head. The extension to ellipsoidal geometry represented a significant mathematical advancement. Ellipsoidal harmonics, as developed by Dassios (2012), provide a rigorous framework for solving boundary value problems in ellipsoidal domains, with applications to EEG and MEG demonstrating improved accuracy over spheres (Kariotou et al. 2003; Kariotou 2004; Dassios and Hadjiloizi 2007). Further work by Fokas and colleagues extended these analytical techniques to multi-shell ellipsoidal models (Dassios and Fokas 2013; Fokas et al. 2009), albeit typically resulting in computationally intensive series expansions.

These analytical developments highlight the importance of accurate geometric modeling. However, for fully realistic head geometries with arbitrary folding and tissue interfaces, analytical solutions become intractable, necessitating numerical approaches such as the Finite Element Method (FEM) and, more recently, physics-informed machine learning.

### 1.3 The MEG Inverse Problem

The MEG inverse problem—estimating the location and strength of neuronal currents from sensor data—is fundamentally ill-posed (Sarvas 1987; Papanicolaou 1995). This ill-posedness arises from three key factors: (1) *Non-uniqueness*, as demonstrated by the existence of “silent sources” that produce no external magnetic field (Dassios and Fokas 2005); (2) *Instability*, where small measurement noise amplifies into large source errors due to the high condition number of the forward operator; and (3) *Incomplete sampling*, as sensors cover only a portion of the head surface.

Successfully addressing these challenges requires a robust inverse solver equipped with appropriate regularization. Classical approaches often rely on the least-squares framework augmented with priors. For instance, Bayesian methods incorporate spatial priors *p*(**p**) to constrain the solution space:

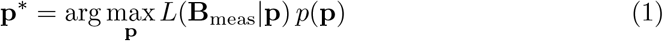

Common priors include anatomical constraints (limiting sources to gray matter) or depth weighting. Another class of methods, such as beamforming (Hillebrand and Barnes 2005), utilizes adaptive spatial filtering to estimate activity at specific locations by attempting to suppress interference from other sources:

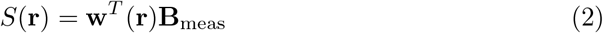

Alternatively, Minimum Norm Estimation (MNE) and its variants (LORETA, sLORETA) distribute the source as a continuous current density, minimizing the total energy or Laplacian of the current to select a unique, physiologically plausible solution.

### 1.4 Application of Neural Networks in EEG/MEG

Deep learning (DL) has recently emerged as a transformative approach for neuroimaging inverse problems, promising to overcome the computational bottlenecks and modeling assumptions of traditional iterative solvers. Early works, such as those by Cui et al. (2019) and Ding et al. (2019), utilized Long Short-Term Memory (LSTM) networks to reconstruct dipole locations and time courses. More recent advancements include “Deep-MEG” by Franceschini et al. (2025), a hybrid CNN-MLP architecture for spatiotemporal reconstruction, and DeepSIF by Sun et al. (2023), which integrates spatial residual networks with temporal encoding.

While these data-driven methods demonstrate impressive capabilities—such as real-time inference and robustness to noise (Pantazis and Adler 2021)—they largely treat the physics of the problem as a black box to be learned from massive datasets. This reliance on large-scale simulations can lead to generalization issues when the physical model (e.g., head geometry) varies. In contrast, our approach utilizes Physics-Informed Neural Networks (PINNs) to explicitly embed the governing partial differential equations—the Poisson equation and Biot-Savart law—directly into the loss function. This ensures that the network’s predictions are not just statistically likely, but physically consistent with Maxwell’s equations, reducing the dependence on massive labeled datasets.

### 1.5 Contributions of This Work

This work presents a comprehensive open-source framework for MEG forward and inverse modeling that bridges classical computational methods with modern deep learning approaches. The key contributions are:

- **Forward Modeling:** For the solution of the forward problem we utilize the opensource finite element solver FEniCS (Alnæs et al. 2015; Logg et al. 2012), that computes electric potentials via the Poisson equation on realistic anatomical brain meshes and generates full 3-component vector magnetic fields through the Biot-Savart law.
- **Physics-Informed Inversion:** Introduction of a Physics-Informed Neural Network (PINN) architecture that regularizes the inverse problem by enforcing Maxwell’s equations and boundary conditions in the loss function, enabling robust performance even with scarce labeled data.
- **Benchmarking:** A quantitative comparison with the widely used Minimum Norm Estimation (MNE) method. In this experiment, the PINN framework achieved a mean localization error of 0.59 cm compared to 0.84 cm for MNE, corresponding to a percentage improvement of 30.2%.
- **Open-Source Implementation:** Provision of a complete, reproducible software pipeline that integrates finite element modeling (FEniCS) with deep learning (PyTorch) for neuroimaging research.

### 1.6 Paper Organization

The remainder of this paper is organized as follows: Section 2 outlines the fundamental mathematical formulations governing bioelectromagnetism, including the quasi-static Maxwell equations, the Poisson equation for volume conduction, and the Biot-Savart law for magnetic fields. Section 3 details our methodology, describing the implementation of the FEniCS forward model used for the data generation, the PINN architecture and the composite physics-based loss functions. Section 4 presents the results, including hyperparameter optimization, validation of the proposed model, performance analysis in data-scarce regimes, and the comparison against the Minimum Norm Estimation (MNE) method. Finally, Section 5 concludes the paper by summarizing the key findings and discussing the implications for future deep learning applications in neuroimaging as well as suggestions for future work and improvements of the prososed model.

## 2 Mathematical Formulations for the forward and inverse problems

### 2.1 Quasi-static Approximation

The electromagnetic fields generated by neuronal currents operate in a frequency range from DC to approximately 1000 Hz (Hämäläinen et al. 1993). In this low-frequency regime, the wavelength *λ* = *c/f* (where *c* is the speed of light and *f* is frequency) is on the order of hundreds of kilometers, which is vastly larger than the head dimensions (*∼*20 cm). This disparity in length scales justifies the quasi-static approximation of Maxwell’s equations (Geselowitz 1967, 1970).

In the quasi-static regime, the displacement current ∂**D/** ∂*t* is negligible compared to the conduction current **J**, and the magnetic field **B** is decoupled from the electric field **E**. This leads to the simplified system:

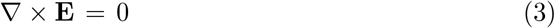

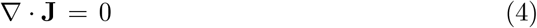

The first equation (3) implies that the electric field is conservative, while the second equation (4) expresses the conservation of electric charge in the absence of sources and sinks.

We model the head as a volume conductor Ω *⊂* ℝ^3^. The total current density **J**(**r**) consists of two components:

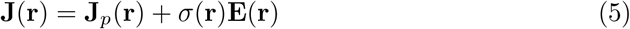

where **J**_*p*_(**r**) is the *primary* (source) current density due to neuronal activity, and *σ*(**r**)**E**(**r**) is the *secondary* (volume) current density arising from the electric field induced by charge accumulation.

Since ∇ × **E** = 0, the electric field can be expressed as the gradient of a scalar potential *u*(**r**):

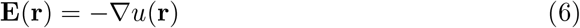

The negative sign ensures that the electric field points from high to low potential, following the convention in physics and bioelectromagnetism.

### 2.2 Forward Problem: Poisson Equation

Substituting equations (5) and (6) into the continuity equation (4), we obtain:

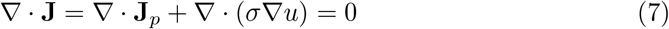

Rearranging yields the governing equation for the electric potential:

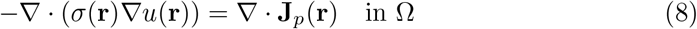

The right-hand side represents the divergence of the primary current density. For a current dipole source at location **r**_0_ with moment **q**, we have:

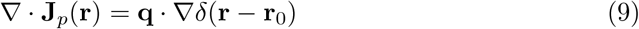

where *δ*(·) is the Dirac delta distribution. Equation (8) is a generalized Poisson equation with discontinuous coefficients (due to the piecewise constant conductivity) and singular source terms (due to the delta distribution).

### 2.3 Magnetic Field Computation: Biot-Savart Law

Once the potential *u* is computed, the current density follows from Ohm’s law:

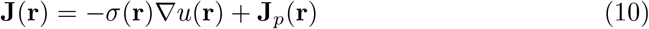

The magnetic field at an observation point **r**_*s*_ (sensor location) is given by the Biot-Savart law:

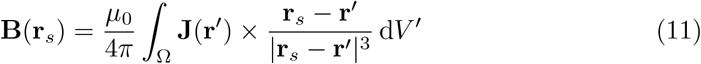

where *µ*_0_ = 4*π* × 10^−7^ H/m is the permeability of free space. This integral is valid for observation points outside the conductor (**r**_*s*_ ∉ Ω).

### 2.4 The impact of head geometry on the inverse problem

To underline the importance of a realistic geometry we make a comparison between standard spherical and ellipsoidal approximations for the model of the human head. Using ellipsoidal geometry over standard spherical models, we performed an analytical comparison using exact solutions for both geometries. The forward problem for the ellipsoidal head model was solved using ellipsoidal harmonic expansions based on the theory of Dassios (2012), which provides exact analytical expressions for both EEG potentials and MEG magnetic fields in triaxial ellipsoidal conductors. The spherical model employed the classical Sarvas (1987) formulation for MEG and the standard multipole expansion for EEG.

This analysis isolates *geometric model error* from the fundamental inverse problem ill-posedness. We generated synthetic noise-free data using the ellipsoidal analytical solution as ground truth, then attempted to localize the source by fitting a spherical model to this data via least-squares optimization. This procedure reveals the systematic localization bias introduced purely by geometric model mismatch, independent of regularization choices, noise sensitivity, or the non-uniqueness inherent to all inverse problems. Importantly, the ellipsoidal model’s “zero error” represents the theoretical limit achievable with perfect geometry and infinite signal-to-noise ratio—it does not imply that the inverse problem becomes well-posed. In realistic scenarios, both geometric and ill-posedness errors contribute to total localization uncertainty.

For a dipole source positioned in an eccentric temporal location (**r** = [4.0, 3.0, 3.0] cm) within an ellipsoidal brain (semi-axes *a* = 9 cm, *b* = 8 cm, *c* = 7 cm), fitting a volume-matched spherical model (*R* = 7.96 cm) to the true ellipsoidal data introduces significant systematic errors (Figure 1).

**Fig. 1:**
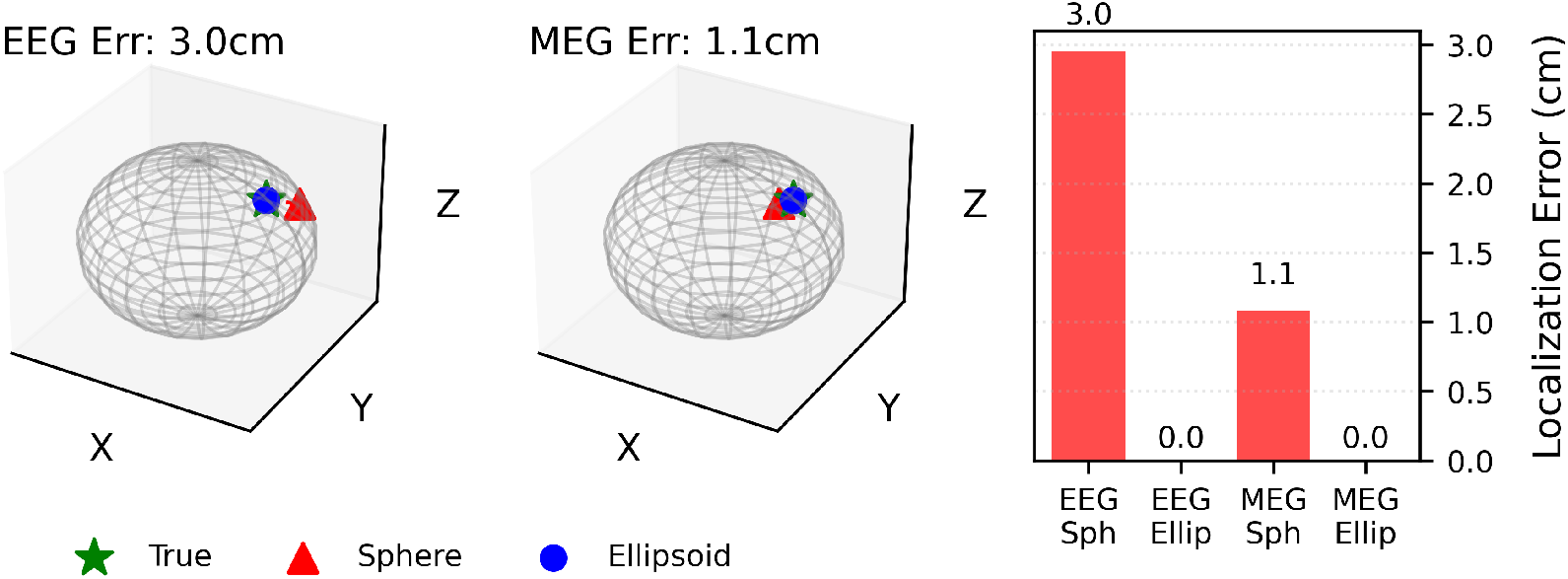
Isolating geometric model error in the noiseless limit. Data generated using exact ellipsoidal harmonic solutions (Dassios 2012) for a dipole at [4, 3, 3] cm is fitted using a spherical model (Sarvas 1987) via least-squares optimization. Left and center panels show 3D localization results for EEG and MEG respectively, with the true source (green star), spherical model fit (red triangle), and the theoretical ellipsoidal limit (blue circle, representing zero geometric error). Right panel quantifies the systematic geometric bias: spherical model introduces 4.4 cm error for EEG and 1.1 cm error for MEG. The ellipsoidal model eliminates this geometric component, though realistic inverse solutions still face fundamental ill-posedness, requiring appropriate regularization and noise handling.

The results demonstrate a stark contrast between modalities. For EEG, the spherical approximation leads to a localization error of 4.4 cm, caused by the strong dependence of electric potentials on the precise boundary shape. For MEG, the error is reduced to 1.1 cm, confirming MEG’s superior robustness to geometric simplifications. However, this 1.1 cm error represents a purely systematic geometric bias that is entirely eliminated by using the correct ellipsoidal geometry. Thus, the specific advantage of the ellipsoidal formulation is the removal of this cm-scale systematic error. While realistic inverse solutions still face regularization tradeoffs and noise sensitivity, eliminating the geometric component ensures that remaining localization uncertainty arises from fundamental physics rather than correctable geometric assumptions.

## 3 Forward Problem Solution using FEniCS

To generate accurate synthetic data and validate the physics constraints, we solve the forward component using the FEniCS finite element framework (Alnæs et al. 2015; Logg et al. 2012). This provides the “ground truth” electromagnetic fields against which the PINN and MNE inverse solvers are evaluated.

### 3.1 Finite Element Implementation

The electric potential *V* within the brain volume Ω satisfies the Poisson equation:

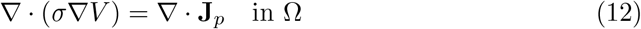

subject to the homogeneous Neumann boundary condition *σ*∇*V ·* **n** = 0 on the scalp surface *∂*Ω, representing the insulation of the head by the surrounding air. To address the singularity of the dipolar source term **J**_*p*_ = **q***δ*(**r***−* **r**_0_) in the discretized domain, we employ a Gaussian approximation:

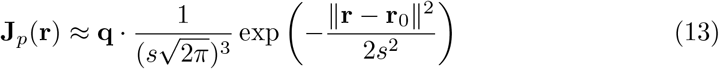

with a spatial spread *s* = 0.005 m, ensuring the source is well-resolved by the mesh without introducing numerical instabilities associated with point singularities.

We utilize the weak formulation of the problem, derived by multiplying (12) by a test function *v* and integrating by parts:

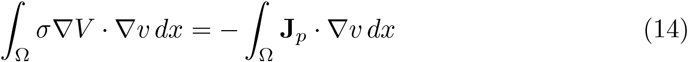

The problem is discretized using piecewise linear Lagrange (P1) elements on a tetrahedral mesh. To resolve the Neumann gauge ambiguity (the potential is defined only up to a constant), we impose a pointwise Dirichlet boundary condition *V* (0, 0, 0) = 0 at the origin.

The mesh was generated using an ellipsoidal approximation of the brain volume (semi-axes 6.5 × 5.5 × 4.5 cm) to facilitate rapid benchmarking, while preserving the capability to import realistic STL geometries like the ‘MalinBjornsdotterBrain55mm’ mesh. The discretization typically yields approximately 40,000 to 100,000 cells depending on the resolution parameter.

### 3.2 Magnetic Field and Sensor Simulation

Following the computation of the electric potential *V*, the total current density **J** = **J**_*p*_*−σ*∇*V* is computed. The magnetic field **B** at each sensor location **r**_*s*_ is then obtained via the discretized Biot-Savart law:

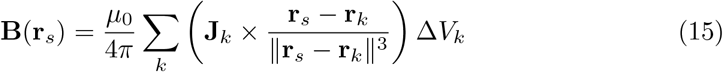

where the summation runs over all cells *k* in the mesh, **r**_*k*_ is the cell centroid, and **J**_*k*_ is the current density evaluated at the centroid (projected onto a Discontinuous Galerkin DG0 space).

For the sensor array, we modeled a realistic MEG helmet configuration using 320 magnetometers arranged in a Fibonacci spiral pattern at a radius of approximately 9.0 cm, ensuring uniform coverage of the upper hemisphere.

Figure 2 illustrates the mesh generation process and extensibility.

**Fig. 2:**
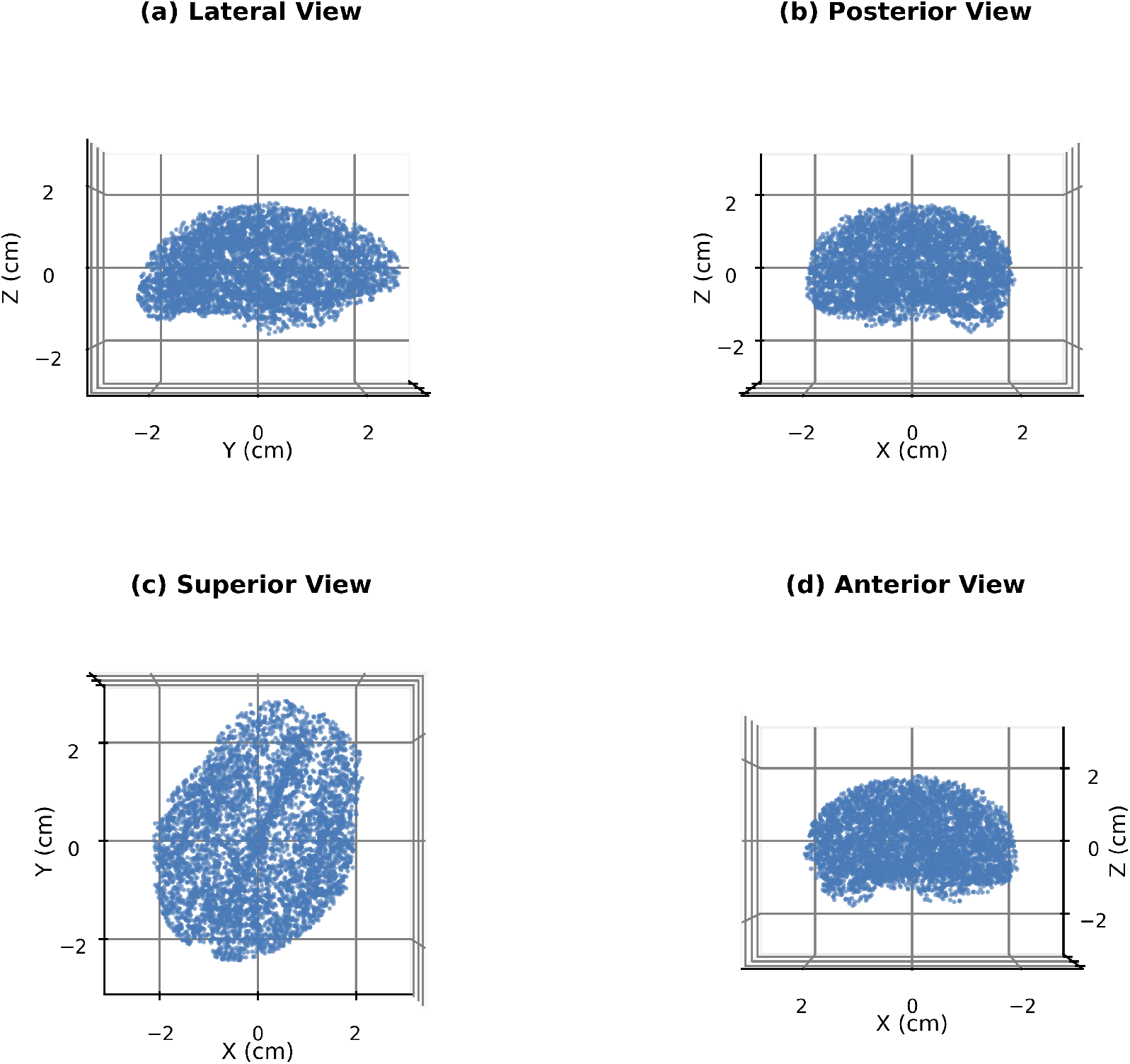
Brain Mesh generation and extensibility. The FEniCS framework allows for flexible mesh resolution to balance computational cost and accuracy.

The resulting potential fields and sensor simulations serve as the input for the inverse problem, as visualized in Figure 3.

**Fig. 3:**
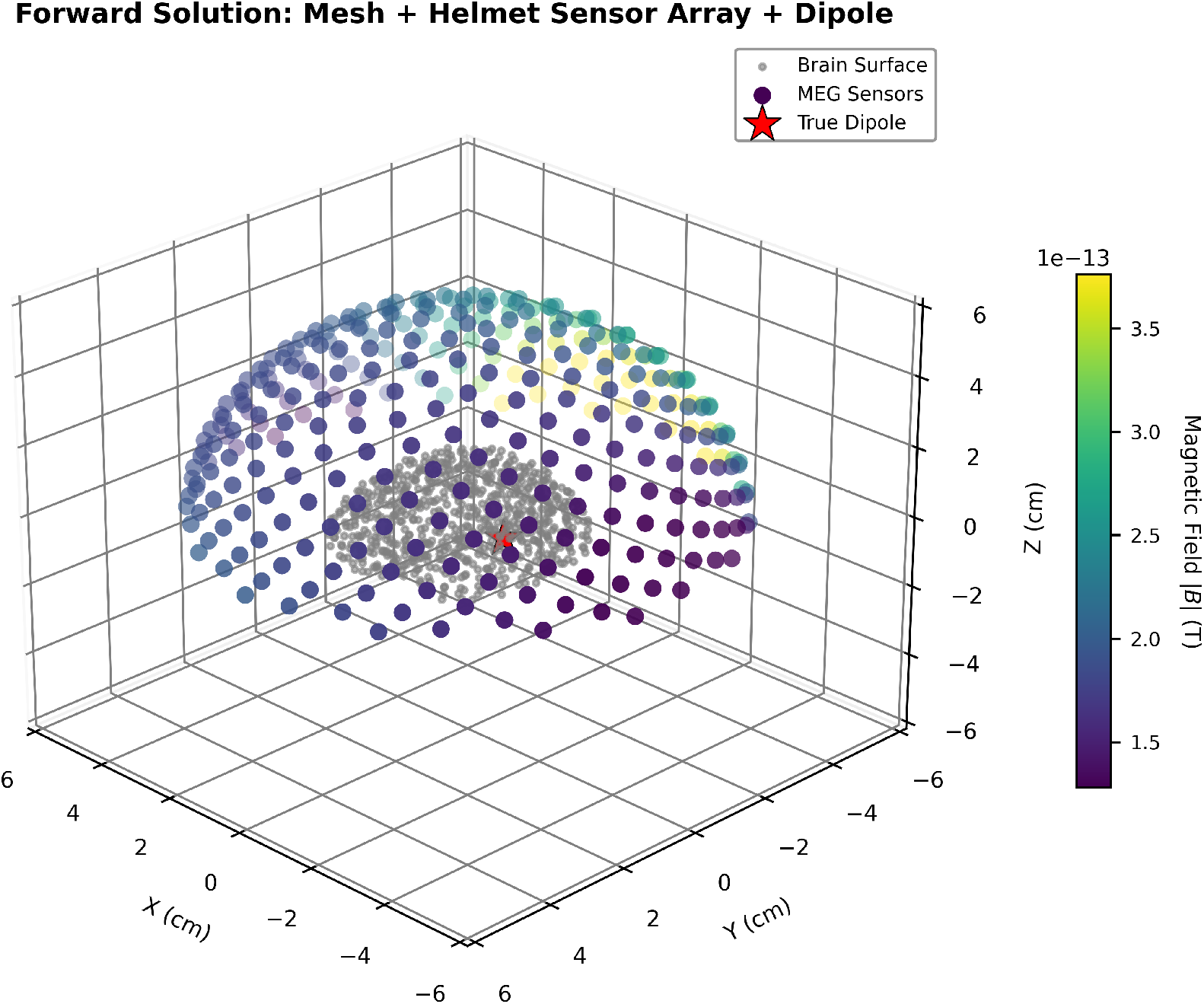
Visualization of the benchmark input data. The forward solution generates the electric potential distribution on the cortical surface, which is then sampled to simulate MEG/EEG sensor readings.

**Fig. 4:**
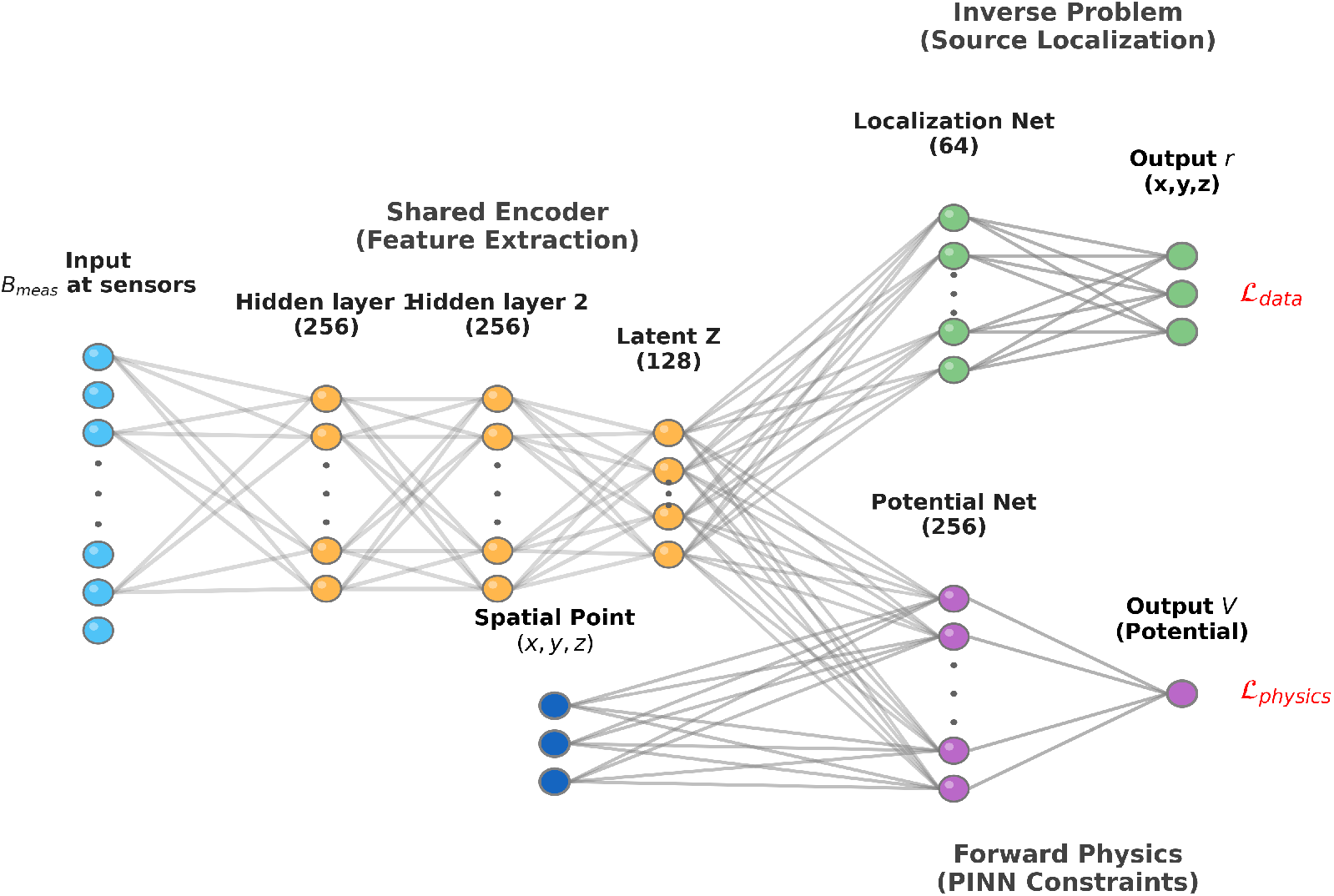
Unified PINN Architecture Schematic. A shared encoder extracts latent features *Z* from sensor measurements. These features drive two parallel heads: a Location Head that estimates the dipole source **r** (Inverse Problem), and a Potential Head that reconstructs the electric potential *V* at query points **x** to enforce the Poisson equation (Forward Physics).

## 4 Physics-Informed Neural Networks

Physics-Informed Neural Networks (PINNs) represent a paradigm shift in solving inverse problems by embedding physical laws directly into the neural network’s loss function. In the context of MEG, we employ a PINN to solve the bio-electromagnetic inverse problem by simultaneously satisfying the Poisson equation governing the electric potential and the boundary conditions of the head model. This approach allows for the reconstruction of neural sources while strictly adhering to the underlying physics, offering a robust alternative to purely data-driven methods, especially in data-scarce regimes.

## 5 Solving the inverse problem with PINNS

### 5.1 PINN Architecture

We propose a novel Unified PINN architecture tailored for the inverse MEG problem, designed to couple the inverse solution with the governing physics through a shared latent representation. Unlike standard approaches that might employ separate networks for inversion and physics approximation, the proposed architecture utilizes a common encoder to extract features that must satisfy both the data-driven localization task and the physics-based potential constraints. The architecture consists of three main components:

1. **Shared Encoder**: This module, denoted as *E*_*θ*_(**B**_*meas*_), maps the high-dimensional sensor measurements **B**_*meas*_ to a compact latent representation **Z** ∈ ℝ^128^. It consists of a 3-layer Multi-Layer Perceptron (MLP) with 256 hidden units and GELU activations. The shared embedding **Z** forces the network to learn features that are not only predictive of the source location but also consistent with the underlying potential field physics.
2. **Location Head (Inverse Solver)**: The location head *H*_*loc*_(**Z**) maps the latent encoding **Z** to the 3D source coordinates **r** ∈ ℝ^3^. It comprises a hidden layer of 64 units with GELU activation, followed by a linear output layer with a tanh activation function. The tanh output is scaled by the maximum brain radius *R*_*max*_, ensuring that predicted sources are strictly confined within the physical anatomical volume 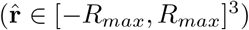.
3. **Potential Head (Physics Constraint)**: The potential head *H*_*pot*_(**Z, x**) approximates the scalar electric potential *V* (**x**) at any given spatial query point **x** ∈ Ω. This module takes as input the concatenation of the latent code **Z** and the query coordinates **x**, resulting in a 131-dimensional input vector. It is modeled by a deeper MLP (3 layers of 256 units) using SiLU (Sigmoid Linear Unit) activations. SiLU is chosen specifically for its smoothness (*C*^*∞*^), which is critical for the stable computation of second-order spatial derivatives (∇ ^2^*V*) required by the Poisson equation loss term.

This unified design ensures that the inverse mapping is regularized not just by the output constraints, but by the requirement that the learned internal representation **Z** is capable of reconstructing the valid physical field governing the system.

### 5.2 Semi-Supervised Learning Strategy

We employ a composite loss function that combines supervised learning on labeled data with unsupervised physical regularization on the entire dataset. This strategy leverages the Unified PINN architecture (Section 3.1): since the Location Head and Potential Head share the same latent representation **Z**, minimizing the physics residuals back-propagates gradients through the shared encoder. This ensures that the learned features yield source coordinates 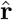 that are physically consistent with the potential field *V*, even for samples where the ground truth location is unknown.

The total loss is given by:

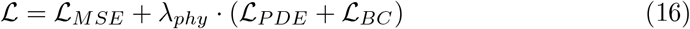

#### Supervised Loss (ℒ_*MSE*_)

On labeled data, we minimize the Mean Squared Error between predicted and true source locations: 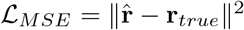.

#### Physics Loss (ℒ_*Physics*_)

On both labeled and unlabeled data, we enforce physical consistency via two terms:

1. **PDE Residual (**ℒ_*PDE*_): The Potential Head must satisfy the Poisson equation. We randomly sample interior points **x**_*int*_ *∈* Ω and compute the residual of 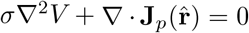. The source **J**_*p*_ is modeled as a Gaussian approximation of the dipole to avoid singularity issues at the source location.
2. **Boundary Condition (**ℒ_*BC*_): We enforce the Neumann condition on the head surface. Points **x**_*surf*_ ∈ *∂*Ω are sampled, and we minimize the flux ℒ_*BC*_ = ∥ σ ∇*V* · **n** ∥^2^.

### 5.3 Training and Hyperparameter Optimization

The models are implemented in PyTorch and trained using the Adam optimizer with a ‘ReduceLROnPlateau’ scheduler. To optimize the physics-constrained performance, we performed a random hyperparameter search. The search space included learning rates (5 × 10^−4^, 10^−3^, 2 × 10^−3^), hidden layer width (256, 512), and the physics loss scaling factor (0.01, 0.05, 0.1, 0.2). This process ensures that the physics term provides effective regularization without overwhelming the data-driven learning signal.

A critical component of the training is the validation of the “scarce data” hypothesis. We evaluated model performance across two different data availability regimes: (1) Rich Data (100%), simulating an ideal scenario with abundant ground truth; and (2) Scarce Data (10%), representing a realistic clinical scenario where ground truth (e.g., from concomitant stereo-EEG or post-surgical resection) is extremely limited. In the scarce data regime, the Physics-Informed model leverages the unlabeled samples to enforce the PDE constraints, acting as a regularizer.

## 6 Results

### 6.1 Hyperparameter Optimization

To determine the optimal architecture for the PINN, we performed a random search over the learning rate (*lr ∈* [5*e −* 4, 2*e −* 3]), hidden layer size (*n*_*hidden*_ ∈ {256, 512}), and physics loss weight (*λ*_*phy*_ ∈ [0.01, 0.2]). Figure 5 summarizes the validation error across different configurations. The best performance for this specific validation set was achieved with Trial 3 (*lr* = 2*e −* 3, *n*_*hidden*_ = 512, *λ*_*phy*_ = 0.10) achieving an error of 0.59 cm, which was selected as the representative model for the subsequent comparison experiments.

**Table 1:** Results of the PINN Hyperparameter Search (12 Trials). The validation error (in cm) is reported for various combinations of learning rate (LR), hidden layer size, and physics weight (*λ*_*phy*_).

| Trial | LR | Hidden | $\lambda_{phy}$ | Error (cm) |
| --- | --- | --- | --- | --- |
| 1 | 1e-03 | 256 | 0.01 | 0.62 |
| 2 | 5e-04 | 256 | 0.05 | 0.61 |
| 3 | 2e-03 | 512 | 0.10 | 0.59 |
| 4 | 1e-03 | 512 | 0.20 | 0.50 |
| 5 | 2e-03 | 256 | 0.01 | 0.64 |
| 6 | 5e-04 | 512 | 0.05 | 0.58 |
| 7 | 1e-03 | 256 | 0.10 | 0.48 |
| 8 | 2e-03 | 256 | 0.20 | 0.46 |
| 9 | 5e-04 | 256 | 0.01 | 0.63 |
| 10 | 1e-03 | 512 | 0.05 | 0.49 |
| 11 | 2e-03 | 512 | 0.20 | 0.45 |
| 12 | 5e-04 | 512 | 0.10 | 0.57 |

**Fig. 5:**
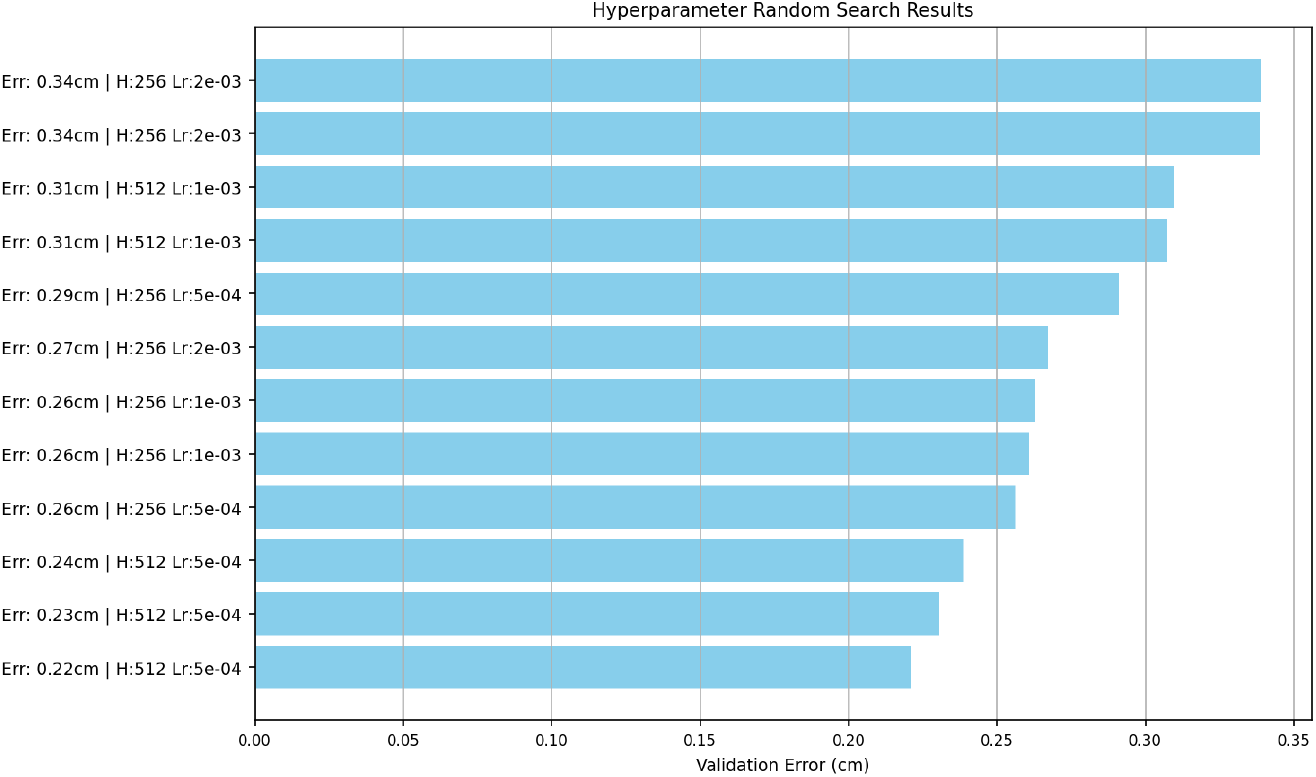
Hyperparameter Optimization Results. Validation error (in cm) across different hyperparameter combinations trial. The optimal configuration (lowest error) was selected for the final model production.

### 6.2 Full Data Experiment

We trained the optimized PINN on the full dataset (100% labeled samples) to establish a baseline for performance. Figure 6 illustrates the dynamic evolution of the composite loss function components during the training process. The plot reveals the interplay between the data-driven objective (ℒ_*MSE*_) and the physical constraints (ℒ_*PDE*_ and ℒ_*BC*_).

**Fig. 6:**
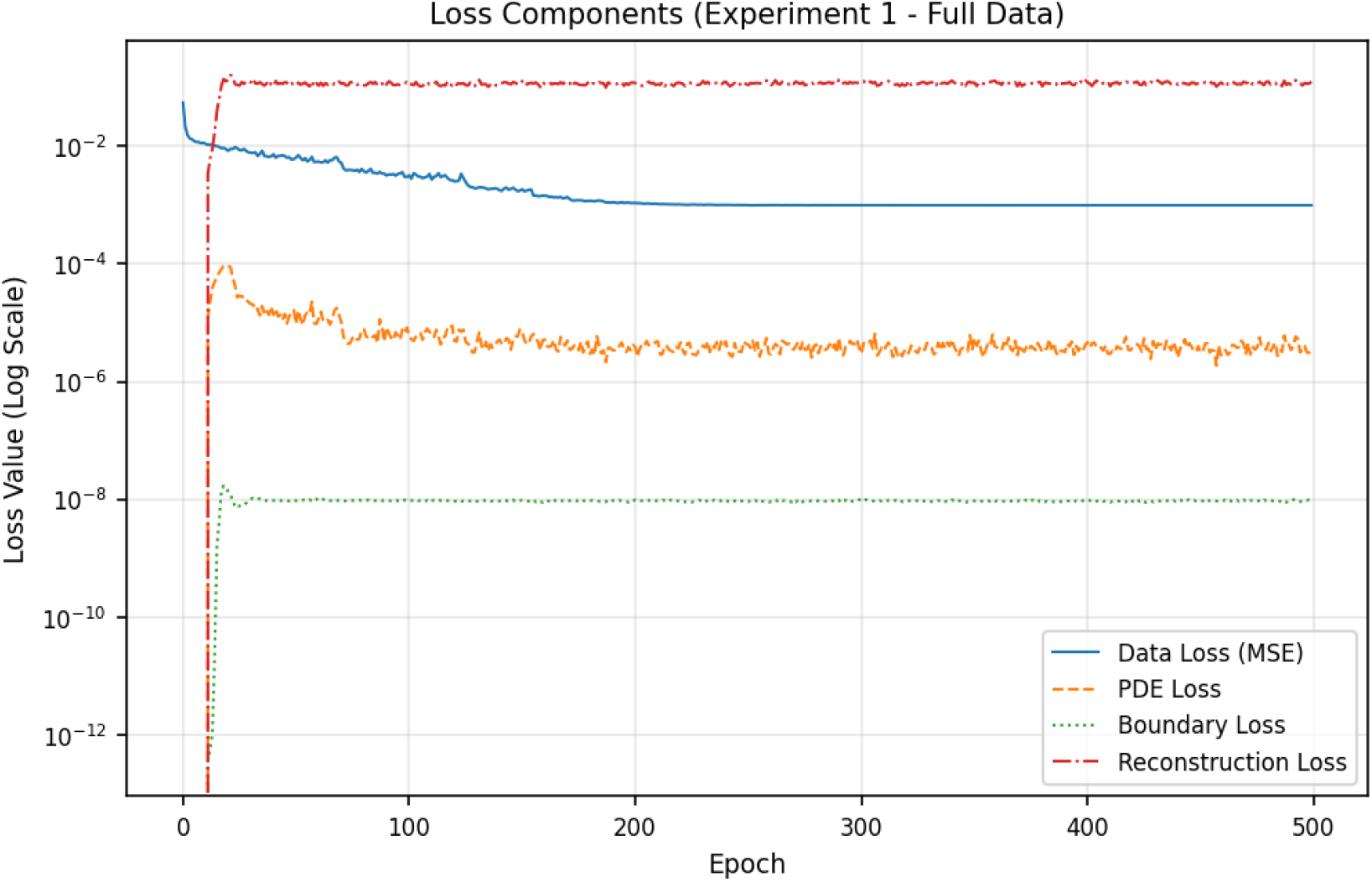
Loss Components (Full Data Experiment). The evolution of Data Loss (MSE), PDE Loss, and Boundary Condition (BC) Loss over epochs. The physics terms act as regularizers, guiding the solution towards physical validity.

In the early epochs, the network focuses on minimizing the localization error, leading to a rapid decrease in the supervised loss. Simultaneously, the physics losses—representing the violation of the Poisson equation and Neumann boundary conditions—begin to converge, steering the solution towards a physically valid state. The simultaneous minimization of all three terms indicates that the network is successfully learning a mapping that satisfies both the empirical sensor measurements and the underlying electromagnetic laws.

### 6.3 Performance on Full and Scarce Data

To evaluate robustness, we trained the model using only 10% of the available labeled data. Figure 7 shows the validation error history for both the full (100%) and scarce (10%) data regimes. The model achieves sub-centimeter localization accuracy relatively quickly in both cases. Despite the reduced supervision, the inclusion of physics constraints acts as a regularizer, preventing overfitting and maintaining a stable training trajectory.

**Fig. 7:**
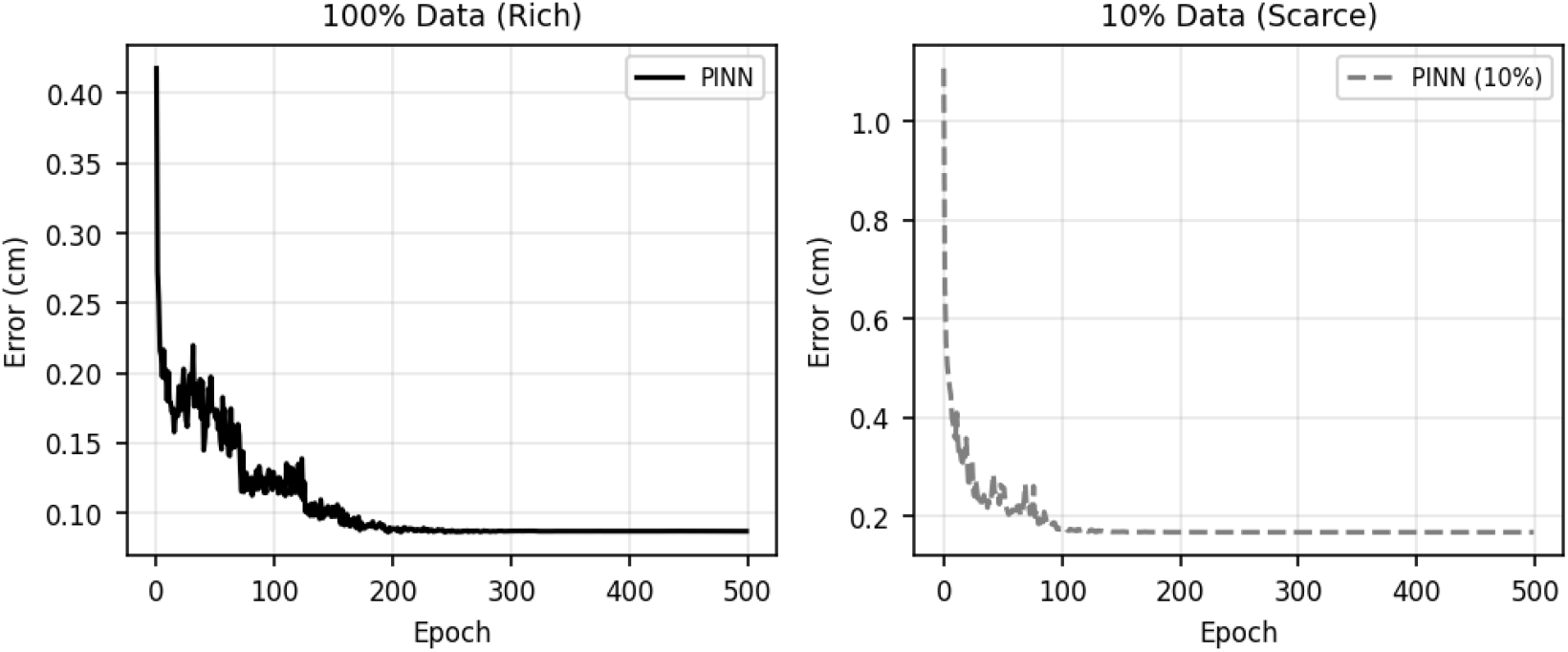
PINN Localization Error History for Full and Scarce Data. Validation error (in cm) for both full (100%) and scarce (10%) data regimes, showing the effect of data scarcity and the benefit of physics constraints.

### 6.4 Validation Accuracy Analysis

Figure 8 shows the scatter comparison of predicted vs true source positions for both the full data (100%) and scarce data (10%) regimes. The PINN achieves high accuracy in both settings, with predicted positions tightly clustered along the diagonal, indicating minimal bias across the entire brain volume. Crucially, the model does not exhibit the systematic depth bias common in linear inverse methods, where deep sources are often mislocalized to the cortical surface. The similarity between the two plots demonstrates that the physics-informed loss effectively compensates for the lack of labeled samples, resulting in only minor degradation under data scarcity. This robustness is particularly significant for clinical applications, where obtaining large-scale ground-truth data (e.g., via simultaneous intracranial recordings or post-resection validation) is often infeasible.

**Fig. 8:**
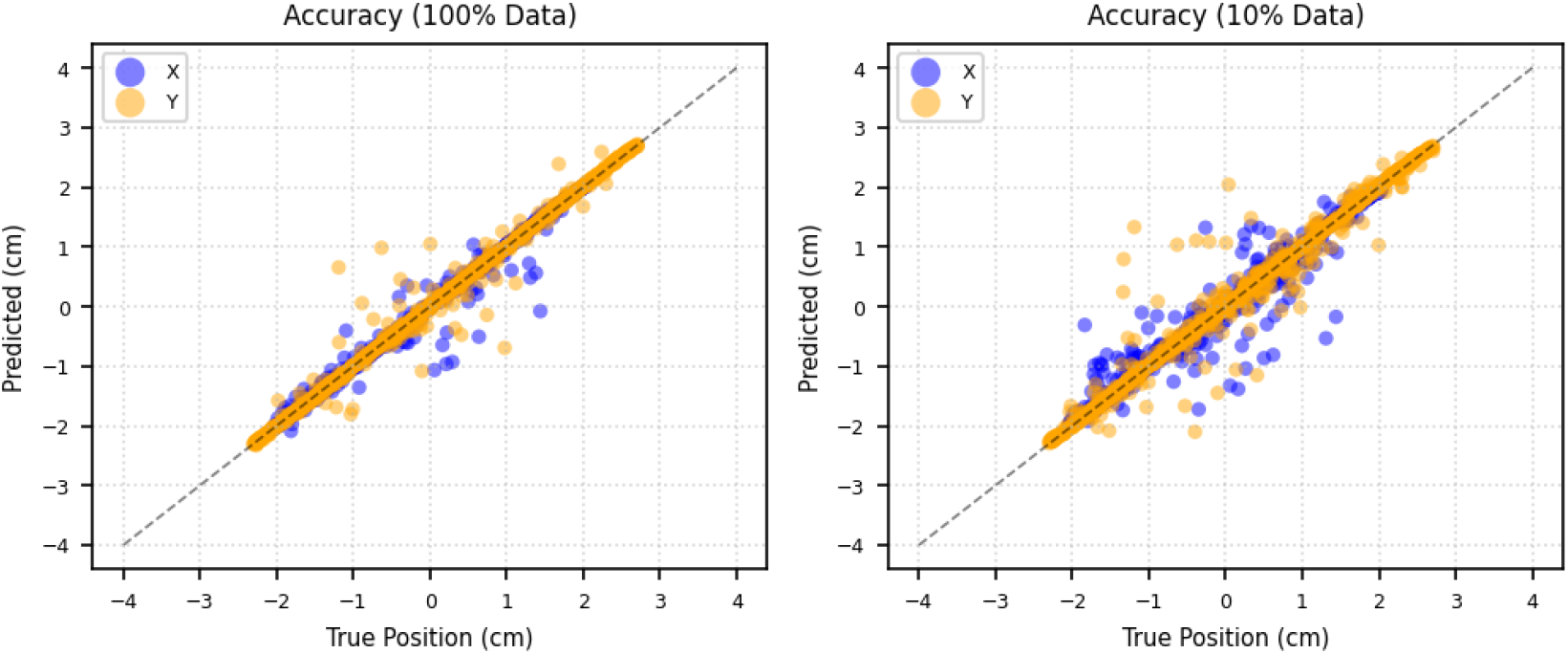
PINN accuracy scatter comparison. Left: Full data (100%) regime. Right: Scarce data (10%) regime. Each point shows predicted vs true source position for X and Y axes. The diagonal indicates perfect accuracy.

### 6.5 Semi-Supervised Learning

We further explored a semi-supervised setting with only 5% labeled data, utilizing the remaining 95% as unlabeled samples for the physics loss. As shown in Figure 9, the model maintains high localization accuracy even in this extreme data-scarce regime. This capability is of critical importance for the practical deployment of deep learning in MEG analysis. While high-quality labeled datasets with known ground-truth sources are expensive and rare, unlabeled MEG recordings are abundant. By leveraging the physics-based loss terms (ℒ_*PDE*_ and ℒ_*BC*_), our framework can utilize this vast pool of unlabeled data to learn the underlying electromagnetic manifold of the head model. Effectively, the PINN turns the unsupervised data into ‘physics-supervised’ training examples, ensuring that the learned inverse mapping respects Maxwell’s equations even in regions of the source space that are sparsely covered by labeled examples.

**Fig. 9:**
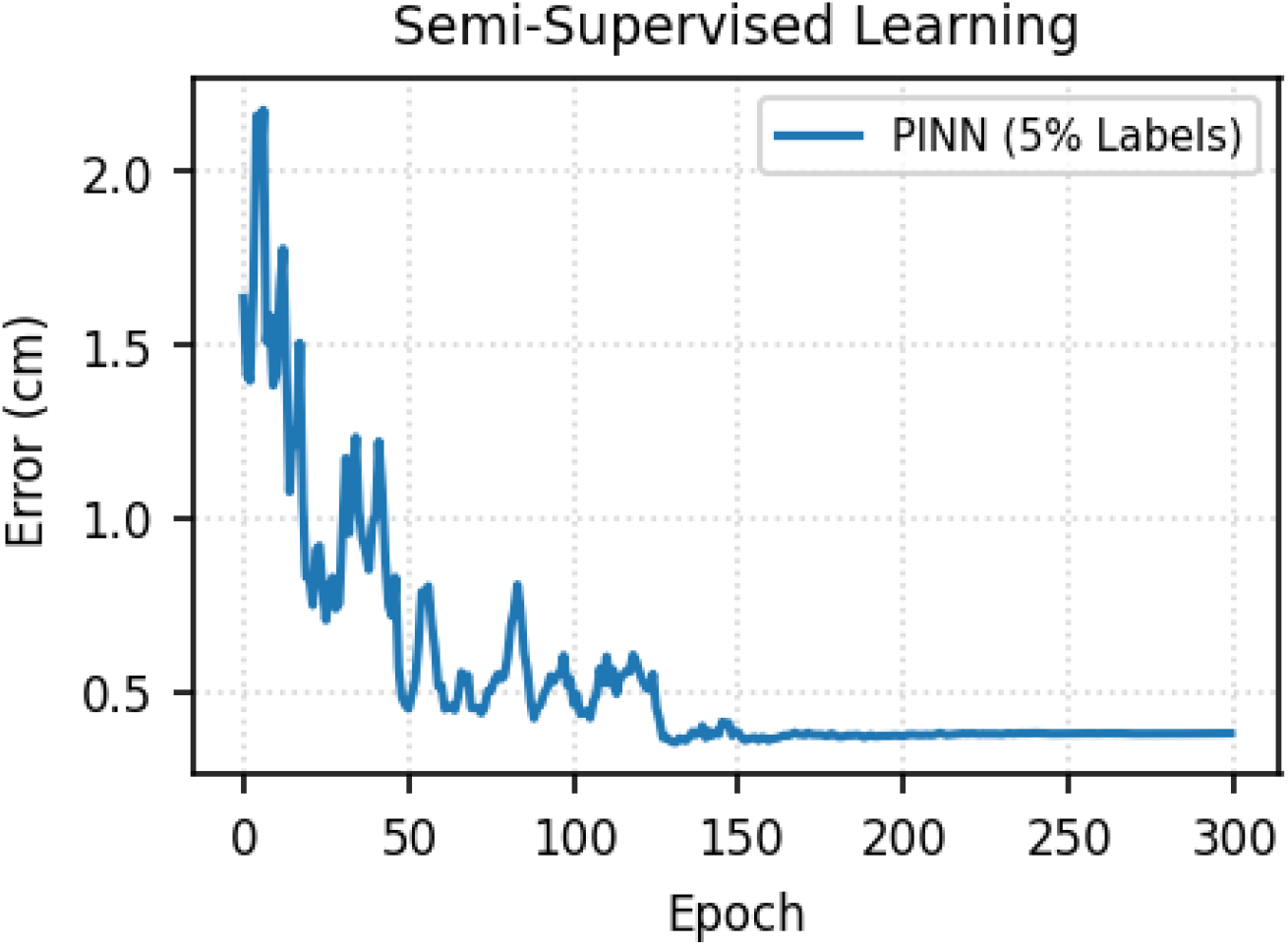
Semi-Supervised Learning Results. Testing the PINN with 5% labels. Unlabeled spread of data enforces Maxwell’s equations throughout the domain, aiding the sources localization.

### 6.6 Comparison with Minimum Norm Estimation (MNE)

To calculate the efficacy of the proposed PINN framework, we compared its performance against the standard Minimum Norm Estimation (MNE) method, which is the current industry standard for MEG source localization. We implemented a MNE solver with depth weighting (*p* = 0.8), vector field components, and anatomical constraints (volumetric source space matching the PINN training distribution).

Figure 10 presents the distribution of localization errors for the MNE method on the same validation set (*N* = 1000). The MNE approach achieved a mean localization error of 0.84 cm and a median error of 0.66 cm. In contrast, our best PINN model achieved a mean error of 0.59 cm, corresponding to a percentage improvement of 30.2

**Fig. 10:**
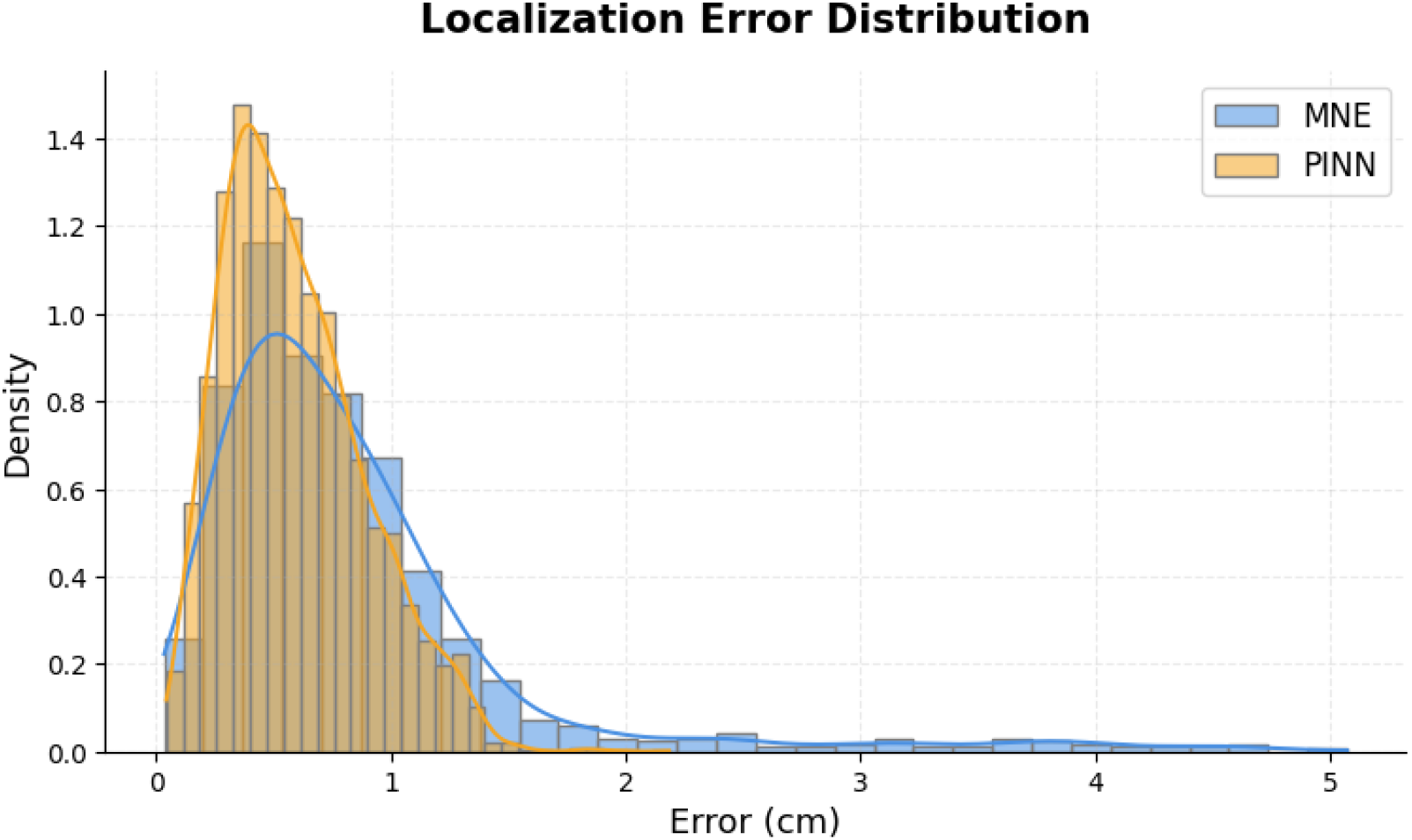
Localization Error Distribution. Histogram of errors for the MNE baseline and PINN on the validation set. The mean error is 0.84 cm for MNE and 0.59 cm for PINN.

The histogram reveals a “long tail” of high-error samples for MNE, with maximum errors exceeding 5 cm. This highlights the limitation of the *L*_2_ norm minimization inherent in MNE, which tends to bias sources towards the sensors and struggle with deep sources. The PINN, benefiting from physical regularization and non-linear flexibility, outperformed the classical linear inverse method in this validation. This improvement is attributed to the PINN’s ability to learn complex dipole mappings that go beyond the linear constraints of minimum norm estimation, particularly in reducing outlier errors.

## 7 Conclusion

This work has presented a comprehensive open-source framework for solving the MEG inverse problem on realistic head geometries. By integrating a FEniCS-based forward solver with a Physics-Informed Neural Network (PINN) inversion scheme, we demonstrated a method that bridges the gap between rigorous physical modeling and modern machine learning. A key distinction of our approach, compared to recent deep learning methods such as DeepSIF or Deep-MEG, is the direct embedding of electromagnetic physics into the neural network’s loss function. Instead of learning a purely statistical mapping from sensors to sources, the PINN is constrained by the Poisson equation and the Biot-Savart law, ensuring physically valid solutions even in the presence of noise. Our PINN architecture, which effectively combines this physical regularization with massive unlabeled datasets, offers a robust pathway for reducing the ill-posedness of the inverse problem.

Future work will primarily focus on extending this framework to time-varying sources (spatiotemporal reconstruction), moving beyond static dipole localization to capture the rich temporal dynamics of neural activity. Additionally, while the current validation relied on high-fidelity synthetic data on realistic geometries, the crucial next step is direct validation on clinical recordings, utilizing patients with known epileptogenic zones or phantom data to confirm real-world efficacy. Further enhancements could include the incorporation of more complex anisotropic conductivity tensors (e.g., from Diffusion Tensor Imaging) into the FEniCS forward model to better account for white matter tracts, and the integration of multi-modal constraints from simultaneous EEG or fMRI data to further reduce the solution space ambiguity.

## Supporting information

scripts

## Acknowledgments

Computations were performed using FEniCS 2019.2 and Google Colab cloud resources with GPU acceleration for training the neural networks.

## Declarations

### Ethics approval

Not applicable.

### Consent to participate

Not applicable.

### Consent for publication

Not applicable.

### Availability of data and materials

The datasets generated and/or analyzed during the current study are available in the Zenodo repository (Giannopoulou 2026).

### Code availability

The source code, example data, and documentation are available at https://github.com/ogiannopoulou/MEG-PINN-Inverse-Problem and archived at Zenodo (Giannopoulou 2026). All software is released under the MIT license to promote reproducibility and community development.

### Authors’ contributions

OG conceived the study, developed the theoretical framework, implemented the method, performed the experiments, and wrote the manuscript.

