## Supplementary material for "Modeling the inverse MEG problem in neuro-imaging using Physics Informed Neural Networks": scripts: fenics_1.py

```
import numpy as np
from dolfin import *
import meshio
import os
import sys

# Configuration
volume_mesh_file = 'MalinBrain_volume.xdmf'
output_filename = 'brain_physics_meta.npz'

# 1. Load 3D Volume Mesh
print(f"Loading volume mesh from {volume_mesh_file}...")
if not os.path.exists(volume_mesh_file):
    print(f"Error: {volume_mesh_file} not found. Please run fenics_2.py first.")
    sys.exit(1)

mesh = Mesh()
with XDMFFile(volume_mesh_file) as infile:
    infile.read(mesh)

print(f"Volume Mesh Loaded: {mesh.num_vertices()} vertices, {mesh.num_cells()} cells")

# 2. Extract Points
# The mesh coordinates are the "interior" points
# PROBLEM: The mesh is too coarse (only ~700 vertices). A PINN needs ~10,000 points.
# SOLUTION: Generate random points inside the convex hull of the mesh.

print("Checking mesh bounds for random point generation...")
coords = mesh.coordinates()
min_b = coords.min(axis=0)
max_b = coords.max(axis=0)

print(f"  Mesh Bounds X: [{min_b[0]:.4f}, {max_b[0]:.4f}]")
print(f"  Mesh Bounds Y: [{min_b[1]:.4f}, {max_b[1]:.4f}]")
print(f"  Mesh Bounds Z: [{min_b[2]:.4f}, {max_b[2]:.4f}]")

# We use the FEniCS Mesh Bounding Box Tree for fast inside/outside checks
bbox_tree = mesh.bounding_box_tree()

# --- NEW LOGIC: Pre-calculate Boundary Hull to Ensure Valid Points ---
print("Extracting boundary for strict containment check...")
bmesh = BoundaryMesh(mesh, "exterior")
surf_pts_check = bmesh.coordinates()
from scipy.spatial import ConvexHull
try:
    hull = ConvexHull(surf_pts_check)
    hull_eqs = hull.equations
    HAS_HULL = True
    print("  Convex Hull built for filtering.")
except Exception as e:
    print(f"  Warning: Could not build Convex Hull ({e}). using only FEniCS bbox check.")
    HAS_HULL = False

def is_strictly_inside(point_arr):
    # point_arr: (N, 3)
    if not HAS_HULL: return np.ones(len(point_arr), dtype=bool)
    # Check: dot(normal, p) + offset <= 1e-5
    # equations: [nx, ny, nz, offset]
    projs = point_arr @ hull_eqs[:, :3].T + hull_eqs[:, 3]
    return np.all(projs <= 1e-4, axis=1)

print("Generating 10,000 strictly interior collocation points...")
interior_pts_list = []
target_count = 10000

# 1. Include Mesh Vertices (Filtered)
mesh_verts = mesh.coordinates()
valid_verts = mesh_verts[is_strictly_inside(mesh_verts)]
print(f"  Mesh Vertices: {len(mesh_verts)} -> Valid: {len(valid_verts)}")

for p in valid_verts:
    interior_pts_list.append(p)

# 2. Fill the rest with random points
np.random.seed(42)
while len(interior_pts_list) < target_count:
    # Batch generation for speed
    batch_size = 2000
    rx = np.random.uniform(min_b[0], max_b[0], batch_size)
    ry = np.random.uniform(min_b[1], max_b[1], batch_size)
    rz = np.random.uniform(min_b[2], max_b[2], batch_size)
    candidates = np.column_stack([rx, ry, rz])
    
    # Filter 1: Convex Hull (Fast Vectorized)
    if HAS_HULL:
        mask_hull = is_strictly_inside(candidates)
        candidates = candidates[mask_hull]
    
    # Filter 2: FEniCS Exact Mesh Check (Slow Loop)
    for p_arr in candidates:
        if len(interior_pts_list) >= target_count: break
        
        p = Point(p_arr[0], p_arr[1], p_arr[2])
        # compute_first_entity_collision returns cell index or -1
        collision = bbox_tree.compute_first_entity_collision(p)
        
        if collision >= 0:
            interior_pts_list.append(p_arr)
    
    if len(interior_pts_list) % 2000 < batch_size: # Approximate print
        print(f"  Collected {len(interior_pts_list)} points...")

interior_pts = np.array(interior_pts_list[:target_count])

# 3. Extract Surface and Normals (Already have bmesh)
# We need the boundary mesh for BC application and surface normals
print("Processing boundary mesh...")
# bmesh already created above
surf_pts = bmesh.coordinates()

# Calculate normals and centers for the boundary mesh cells
print("Computing surface normals and centers...")
normals = []
centers = []
# Iterate over facets (triangles) of the boundary mesh
for i in range(bmesh.num_cells()):
    # Get the cell (triangle)
    cell = Cell(bmesh, i)
    
    # Get Normal (outward facing by default for BoundaryMesh)
    n = cell.cell_normal()
    normals.append([n.x(), n.y(), n.z()])
    
    # Get Midpoint
    m = cell.midpoint()
    centers.append([m.x(), m.y(), m.z()])

surf_normals = np.array(normals)
surf_midpoints = np.array(centers)

print(f"Interior points (Volume Vertices): {len(interior_pts)}")
print(f"Surface points (Boundary Vertices): {len(surf_pts)}")
print(f"Surface elements (Triangles): {bmesh.num_cells()}")
print(f"Surface normals calculated: {len(surf_normals)}")

# 4. Save Data
print(f"Saving to {output_filename}...")
np.savez(output_filename, 
         interior_pts=interior_pts, 
         surface_pts=surf_pts,
         surface_midpoints=surf_midpoints,
         surface_normals=surf_normals)

print("✓ Done. Physics metadata extracted from volumetric mesh.")
```
