## Supplementary material for "Modeling the inverse MEG problem in neuro-imaging using Physics Informed Neural Networks": scripts: fenics_2.py

```
#!/usr/bin/env python3
"""
MEG Training Data Generator
===========================
Generates training data for PINN by solving forward problems with random dipole locations.
"""

import numpy as np
import os
import meshio
from dolfin import *
from mshr import *

# Set FEniCS log level
set_log_level(LogLevel.ERROR)

# ============================================================================
# 1. MESH SETUP - Generate Volumetric Mesh from STL
# ============================================================================
stl_filename = 'MalinBjornsdotterBrain55mm.stl'
volume_mesh_file = 'MalinBrain_volume.xdmf'

def generate_volumetric_mesh_pygalmesh(stl_path, output_path, max_edge_length=0.003):
    """Generate volumetric tetrahedral mesh from STL using pygalmesh"""
    print(f"Generating volumetric mesh from {stl_path} using pygalmesh...")
    
    try:
        import pygalmesh
        
        # Load STL
        mesh_stl = meshio.read(stl_path)
        points = mesh_stl.points / 1000.0  # mm -> m
        center = points.mean(axis=0)
        points = points - center
        
        print(f"  STL vertices: {len(points):,}, triangles: {len(mesh_stl.cells_dict['triangle']):,}")
        print(f"  Center: [{center[0]:.4f}, {center[1]:.4f}, {center[2]:.4f}] m")
        
        # Create temporary STL with centered coordinates
        temp_stl = 'temp_centered_brain.stl'
        mesh_centered = meshio.Mesh(points, [("triangle", mesh_stl.cells_dict['triangle'])])
        meshio.write(temp_stl, mesh_centered)
        
        # Generate volume mesh using pygalmesh
        print(f"  Running tetrahedral mesh generation (max edge: {max_edge_length*100:.2f} cm)...")
        print(f"  This may take 1-3 minutes for the Malin brain mesh...")
        
        # Adjust parameters for better mesh generation
        # max_edge_length is in meters (0.008 = 8mm)
        # We might need slightly finer mesh or different parameters
        mesh_vol = pygalmesh.generate_volume_mesh_from_surface_mesh(
            temp_stl,
            max_edge_size_at_feature_edges=max_edge_length,
            max_cell_circumradius=max_edge_length * 2,
            min_facet_angle=25,
            max_radius_surface_delaunay_ball=max_edge_length,
            verbose=True
        )
        
        print(f"  Raw Generated: {len(mesh_vol.points):,} vertices")
        for cell_type, cells in mesh_vol.cells_dict.items():
            print(f"    {cell_type}: {len(cells):,} cells")
        
        # Filter to keep only tetrahedra for FEniCS
        if 'tetra' in mesh_vol.cells_dict:
            tetra_cells = mesh_vol.cells_dict['tetra']
            print(f"  Extracting {len(tetra_cells):,} tetrahedra for FEniCS...")
            
            # Create a clean mesh with only tetrahedra
            mesh_out = meshio.Mesh(
                points=mesh_vol.points,
                cells=[("tetra", tetra_cells)]
            )
            
            # Save as XDMF for FEniCS
            print(f"  Saving volumetric mesh (tetra only) to {output_path}")
            meshio.write(output_path, mesh_out)
            
            # Additional check: If too few cells, something went wrong
            if len(tetra_cells) < 100:
                print("  Warning: Mesh has too few tetrahedra. Generation might have failed.")
                return False, None
                
            # Cleanup
            if os.path.exists(temp_stl):
                os.remove(temp_stl)
            
            return True, center
        else:
            print("  Error: No tetrahedra generated!")
            return False, None
        
    except ImportError:
        print("  pygalmesh not available")
        return False, None
    except Exception as e:
        print(f"  Volumetric mesh generation failed: {e}")
        import traceback
        traceback.print_exc()
        return False, None

def create_fenics_mesh(stl_path, volume_path, res=24):
    """Load or create volumetric mesh"""
    
    # Try to load existing volumetric mesh
    if os.path.exists(volume_path):
        print(f"Loading existing volumetric mesh: {volume_path}")
        try:
            mesh = Mesh()
            with XDMFFile(volume_path) as infile:
                infile.read(mesh)
            
            coords = mesh.coordinates()
            radii = np.array([
                (coords[:,0].max() - coords[:,0].min()) / 2,
                (coords[:,1].max() - coords[:,1].min()) / 2,
                (coords[:,2].max() - coords[:,2].min()) / 2
            ])
            print(f"  Loaded: {mesh.num_vertices():,} vertices, {mesh.num_cells():,} cells")
            return mesh, radii
        except Exception as e:
            print(f"  Failed to load {volume_path}: {e}")
            if os.path.exists(volume_path):
                os.remove(volume_path)
    
    # Try to generate volumetric mesh from STL using pygalmesh
    if os.path.exists(stl_path):
        success, center = generate_volumetric_mesh_pygalmesh(stl_path, volume_path, max_edge_length=0.003)
        if success:
            return create_fenics_mesh(stl_path, volume_path, res)
    
    # Fallback: Create ellipsoid approximation
    print("Using ellipsoid approximation (fallback)...")
    if os.path.exists(stl_path):
        mesh_in = meshio.read(stl_path)
        points = mesh_in.points / 1000.0
        center = points.mean(axis=0)
        points = points - center
        radii = (points.max(axis=0) - points.min(axis=0)) / 2
    else:
        radii = np.array([0.065, 0.055, 0.045])
    
    print(f"  Creating ellipsoid with radii: [{radii[0]*100:.2f}, {radii[1]*100:.2f}, {radii[2]*100:.2f}] cm")
    mesh = generate_mesh(domain, res)
    return mesh, radii

print("Creating volumetric mesh...")
mesh, radii_m = create_fenics_mesh(stl_filename, volume_mesh_file)
V_space = FunctionSpace(mesh, 'P', 1)
print(f"✓ Mesh ready: {mesh.num_vertices():,} vertices, {mesh.num_cells():,} cells")
print(f"  Radii (cm): [{radii_m[0]*100:.2f}, {radii_m[1]*100:.2f}, {radii_m[2]*100:.2f}]")

# Get valid range for random dipole placement
coords = mesh.coordinates()
min_x, max_x = coords[:,0].min(), coords[:,0].max()
min_y, max_y = coords[:,1].min(), coords[:,1].max()
min_z, max_z = coords[:,2].min(), coords[:,2].max()
print(f"Valid dipole range (m): X=[{min_x:.4f},{max_x:.4f}], Y=[{min_y:.4f},{max_y:.4f}], Z=[{min_z:.4f},{max_z:.4f}]")

# ============================================================================
# 2. SENSOR SETUP (Fibonacci spiral on helmet)
# ============================================================================
n_sensors = 320
helmet_radius = np.max(radii_m) + 0.03
sensors = []
golden_ratio = (1 + np.sqrt(5)) / 2
for i in range(n_sensors):
    t = i / (n_sensors - 1)
    z_rel = t
    radius_at_z = np.sqrt(1 - z_rel**2)
    theta = 2 * np.pi * i / golden_ratio
    sensors.append([helmet_radius * radius_at_z * np.cos(theta),
                    helmet_radius * radius_at_z * np.sin(theta),
                    helmet_radius * z_rel])
sensors = np.array(sensors)
print(f"MEG sensors: {len(sensors)}")

# ============================================================================
# 3. FORWARD SOLVER SETUP
# ============================================================================
sigma = 0.33  # S/m
mu0_4pi = 1e-7

# Pre-calculate cell data for Biot-Savart
mesh_cells = list(cells(mesh))
cell_midpoints = np.array([c.midpoint().array() for c in mesh_cells])
cell_volumes = np.array([c.volume() for c in mesh_cells])

def solve_forward_and_compute_meg(dipole_pos):
    """Solve forward EEG problem and compute MEG field for given dipole position using PointSource approx"""
    
    # 1. Define Dipole as two point sources (Monopole Approximation)
    # Q = I * d. Let d = 2mm.
    # We orient the dipole in X direction for training consistency
    d = 0.002 # 2 mm separation
    Q_mag = 1e-8 # 10 nAm (Stronger signal for training visibility)
    I = Q_mag / d
    
    pos_plus = Point(dipole_pos[0] + d/2, dipole_pos[1], dipole_pos[2])
    pos_minus = Point(dipole_pos[0] - d/2, dipole_pos[1], dipole_pos[2])
    
    # 2. Assemble System
    u = TrialFunction(V_space)
    v = TestFunction(V_space)
    a = sigma * dot(grad(u), grad(v)) * dx
    L = Constant(0) * v * dx # Zero volume source, we use PointSource applied to vector
    
    # Assemble A and b
    A = assemble(a)
    b = assemble(L)
    
    # Apply Point Sources
    # Check if points are in mesh (they should be, but robustness check)
    try:
        PointSource(V_space, pos_plus, I).apply(b)
        PointSource(V_space, pos_minus, -I).apply(b)
    except Exception:
        # If point is outside (e.g. near boundary), ignore or raise
        pass
        
    # Boundary condition (None/Natural for insulated skull/air interface in this simplified model)
    # Reference: Brain boundary grad(u).n = 0 implies no current leaves the brain.
    
    # Solve
    u_sol = Function(V_space)
    solve(A, u_sol.vector(), b)
    
    # 3. Compute current density J = -sigma * grad(u) - J_p
    # Note: total current J_tot = -sigma*grad(u) + J_primary
    # The MEG field is B = B_vol + B_prim
    # B_prim is the field from the primary dipole in infinite medium.
    # B_vol is from volume currents J_vol = -sigma*grad(u).
    
    # A. Volume Currents Contribution (Numerical)
    # Project gradient to DG space (piecewise constant on cells)
    V_dg = VectorFunctionSpace(mesh, 'DG', 0)
    J_vol_field = project(-sigma * grad(u_sol), V_dg)
    
    # Get J values at each cell center
    J_vol_array = J_vol_field.vector().get_local()
    n_cells = len(mesh_cells)
    J_vol_vals = J_vol_array.reshape((n_cells, 3))
    
    # Biot-Savart for Volume Currents
    # B_vol = mu0/4pi * sum( J_vol x r / r^3 * dV )
    B_field = np.zeros((len(sensors), 3))
    
    # Vectorized Biot-Savart
    # Only loop over sensors (320), vectorize over cells (2500)
    for i in range(len(sensors)):
        r_vec = sensors[i] - cell_midpoints # (N_cells, 3)
        r_mag = np.linalg.norm(r_vec, axis=1)
        r_mag_cubed = r_mag**3 + 1e-15
        
        # Cross product J x r
        # J: (N, 3), r: (N, 3) -> cross: (N, 3)
        cross_prod = np.cross(J_vol_vals, r_vec)
        
        # Integration
        contribution = (mu0_4pi * cross_prod / r_mag_cubed[:, np.newaxis]) * cell_volumes[:, np.newaxis]
        B_vol_vec = np.sum(contribution, axis=0) # (3,)
        
        # Add primary contribution (dipole formula)
        # B_prim = mu0/4pi * (Q x r) / r^3
        r_p = sensors[i] - dipole_pos
        r_p_mag = np.linalg.norm(r_p)
        Q_vec = np.array([Q_mag, 0, 0]) # X-oriented
        B_prim_vec = mu0_4pi * np.cross(Q_vec, r_p) / (r_p_mag**3 + 1e-12)
        
        # Total B vector
        B_total = B_vol_vec + B_prim_vec
        
        # Record full vector (Bx, By, Bz) instead of scalar magnitude
        B_field[i] = B_total
    
    return B_field

# ============================================================================
# 4. GENERATE TRAINING DATA
# ============================================================================
n_samples = 5000  # Generate 5000 samples for actual training
print(f"\nGenerating {n_samples} training samples...")

X_data = []  # Magnetic field measurements
Y_data = []  # Dipole locations

np.random.seed(42)  # For reproducibility

# Ensure we pick points INSIDE the new volumetric mesh
# We can use the mesh.bounding_box_tree() but for complex geometries
# it's better to pick from actual mesh cells or use rejection sampling
mesh_bbox_tree = mesh.bounding_box_tree()

count = 0
while count < n_samples:
    # 1. Random point within bounding box
    p_x = np.random.uniform(min_x, max_x)
    p_y = np.random.uniform(min_y, max_y)
    p_z = np.random.uniform(min_z, max_z)
    point = Point(p_x, p_y, p_z)
    
    # 2. Check if inside mesh
    # compute_first_entity_collision returns cell index or -1
    collision = mesh_bbox_tree.compute_first_entity_collision(point)
    
    if collision < 0:
        continue  # Point outside brain, skip
        
    dipole_pos = np.array([p_x, p_y, p_z])
    
    # Solve forward problem
    if count % 50 == 0:
        print(f"Solving sample {count+1}/{n_samples}...")
        
    try:
        B_field = solve_forward_and_compute_meg(dipole_pos)
        X_data.append(B_field)
        Y_data.append(dipole_pos)
        count += 1
    except Exception as e:
        print(f"Solver failed for point {dipole_pos}: {e}")

# Save data
X_data = np.array(X_data)
Y_data = np.array(Y_data)

np.savez('meg_training_data.npz', 
         X=X_data,  # Shape: (n_samples, n_sensors)
         Y=Y_data,  # Shape: (n_samples, 3)
         sensors=sensors)

print(f"\n✓ Saved meg_training_data.npz")
print(f"  X (MEG fields): {X_data.shape}")
print(f"  Y (dipole positions): {Y_data.shape}")
```
