## Supplementary material for "Modeling the inverse MEG problem in neuro-imaging using Physics Informed Neural Networks": scripts: fenics_forward_poisson.py

```
#!/usr/bin/env python3
"""
FEniCS Forward Model Solver for MEG (Poisson Equation)
======================================================
This script solves the forward problem using FEniCS and saves the synthetic data
for use in the PINN benchmark notebook.

Physical Model:
1. Poisson Equation: -sigma * div(grad(V)) = div(Jp)
2. Boundary Condition: sigma * grad(V).n = Jp.n (J_tot.n = 0)
3. Biot-Savart Integration for B field.
"""

import numpy as np
import os
import sys
import meshio

try:
    from dolfin import *
    from mshr import *
    set_log_level(LogLevel.ERROR)
    print("FEniCS detected.")
except ImportError:
    print("Error: FEniCS (dolfin/mshr) not installed.")
    sys.exit(1)

# ============================================================================
# 1. GEOMETRY GENERATION
# ============================================================================

stl_filename = 'MalinBjornsdotterBrain55mm.stl'
output_filename = 'fenics_poisson_forward_data.npz'

def create_fenics_mesh(stl_path, res=24):
    if not os.path.exists(stl_path):
        # Fallback dimensions if STL missing (for testing)
        print(f"Warning: {stl_path} not found. Using default ellipsoid.")
        radii = np.array([0.065, 0.055, 0.045])
        center = np.array([0.0, 0.0, 0.0])
    else:
        print("Loading STL dimensions...")
        mesh_in = meshio.read(stl_path)
        points = mesh_in.points / 1000.0 # mm -> m
        center = points.mean(axis=0)
        points = points - center
        radii = (points.max(axis=0) - points.min(axis=0)) / 2
    
    print(f"Approximating with Ellipsoid: Radii={radii*100} cm")
    domain = Ellipsoid(Point(0,0,0), radii[0], radii[1], radii[2])
    mesh = generate_mesh(domain, res)
    print(f"Mesh Generated: {mesh.num_vertices()} vertices, {mesh.num_cells()} cells")
    
    return mesh, radii, center

mesh, radii_m, center_orig_m = create_fenics_mesh(stl_filename, res=24)
V_space = FunctionSpace(mesh, 'P', 1)

# ============================================================================
# 2. SOLVE POISSON EQUATION
# ============================================================================

sigma = 0.33  # S/m
dipole_pos_rel = np.array([0.01, 0.005, 0.005]) # meters (Inside the small brain)
dipole_moment = np.array([1e-8, 0.0, 0.0])   # 10 nA.m

print("Setting up Variational Problem...")

class GaussianDipole(UserExpression):
    def __init__(self, pos, moment, spread_m, **kwargs):
        self.pos = pos
        self.mom = moment
        self.spread = spread_m  # Spatial spread in meters (renamed to avoid confusion)
        # Normalization to ensure Volume Integral matches Moment
        self.norm_factor = 1.0 / ((self.spread * np.sqrt(2*np.pi))**3)
        super().__init__(**kwargs)
    def eval(self, value, x):
        r = x - self.pos
        dist_sq = np.sum(r**2)
        gauss = np.exp(-dist_sq / (2 * self.spread**2))
        value[:] = self.mom * gauss * self.norm_factor
    def value_shape(self):
        return (3,)

Jp_expr = GaussianDipole(dipole_pos_rel, dipole_moment, spread_m=0.005, degree=1)

u = TrialFunction(V_space)
v = TestFunction(V_space)
a = sigma * inner(grad(u), grad(v)) * dx
# Correct weak form: -sigma*Laplacian(V) = div(Jp) -> sigma*grad(u).grad(v) = -Jp.grad(v)
L = -inner(Jp_expr, grad(v)) * dx  # Note: negative sign is critical! 

# Fix potential at origin (Reference) to handle non-uniqueness
bc = DirichletBC(V_space, Constant(0.0), "near(x[0], 0) && near(x[1], 0) && near(x[2], 0)", method="pointwise")

print("Solving...")
V_sol = Function(V_space)
solve(a == L, V_sol, bc)
print("Potential V computed.")

# ============================================================================
# 3. COMPUTING B FIELD
# ============================================================================

# Sensors (Helmet Configuration - Fibonacci Spiral)
print("Generating Helmet Sensor Configuration...")
n_sensors = 320
# Adjust helmet radius relative to approximated ellipsoid
# We want it about 2-3 cm away from the scalp surface
# Max brain radius ~6.5cm. Helmet ~9.0cm
helmet_radius = np.max(radii_m) + 0.03 

sensors = []
golden_ratio = (1 + np.sqrt(5)) / 2
for i in range(n_sensors):
    # z goes from 0 to 1 (top hemisphere)
    # We want sensors from equator up, so z relative to center from 0 to R
    # Actually, a spiral on a sphere is usually z from -1 to 1. 
    # We want the top hemisphere: z from 0 to 1
    
    t = i / (n_sensors - 1) # 0 to 1
    # vertical position (0 to 1 for hemisphere)
    z_rel = t 
    # Radius at this z slice
    radius_at_z = np.sqrt(1 - z_rel**2)
    theta = 2 * np.pi * i / golden_ratio
    
    x = helmet_radius * radius_at_z * np.cos(theta)
    y = helmet_radius * radius_at_z * np.sin(theta)
    z = helmet_radius * z_rel
    
    # Adjust center if needed, but mesh is centered at 0,0,0
    # Add some offset to z if we want to cover more of the side, 
    # but z_rel=0 puts sensors at z=0 (equator). 
    
    sensors.append([x, y, z])

sensors = np.array(sensors)

# Filter any that might be too close (safety) - though helmet_radius should prevent this
dist_from_center = np.linalg.norm(sensors, axis=1)
valid_mask = dist_from_center > np.max(radii_m) + 0.015
sensors = sensors[valid_mask]

print(f"Sensors generated: {len(sensors)} (Helmet Radius: {helmet_radius:.3f} m)")

# Integrate B field using cell-based quadrature (more accurate)
print("Integrating B field...")
V_vec = VectorFunctionSpace(mesh, 'P', 1)
J_total_proj = project(Jp_expr - sigma * grad(V_sol), V_vec)

# Use DG0 (cell-centered) for more accurate volume integration
DG0 = FunctionSpace(mesh, 'DG', 0)
J_x = project(J_total_proj[0], DG0)
J_y = project(J_total_proj[1], DG0)
J_z = project(J_total_proj[2], DG0)

# Get cell centers and volumes
mesh_coords = []
cell_volumes = []
J_vals = []

for cell in cells(mesh):
    mesh_coords.append(cell.midpoint().array())
    cell_volumes.append(cell.volume())
    # Get J values at cell center
    J_vals.append([J_x(cell.midpoint()), J_y(cell.midpoint()), J_z(cell.midpoint())])

mesh_coords = np.array(mesh_coords)
cell_volumes = np.array(cell_volumes)
J_vals = np.array(J_vals)

print(f"Using {len(mesh_coords)} cells for integration (was {mesh.num_vertices()} vertices)")
total_vol = np.sum(cell_volumes)


B_meas = np.zeros((len(sensors), 3))
mu0 = 4 * np.pi * 1e-7

for i, r_s in enumerate(sensors):
    r_vec = r_s - mesh_coords
    r_mag = np.linalg.norm(r_vec, axis=1, keepdims=True)
    r_mag = np.maximum(r_mag, 1e-6)  # Avoid singularities
    cross_prod = np.cross(J_vals, r_vec)
    # Volume-weighted integration
    integrand = (mu0 / (4 * np.pi)) * cross_prod / (r_mag**3)
    B_meas[i] = np.sum(integrand * cell_volumes[:, None], axis=0)

B_mag = np.linalg.norm(B_meas, axis=1)
print(f"Max B: {np.max(B_mag):.2e} T")

# ============================================================================
# 4. SAVE RESULTS
# ============================================================================

np.savez(output_filename, 
         radii_m=radii_m,
         center_m=center_orig_m,
         dipole_pos=dipole_pos_rel,
         dipole_moment=dipole_moment,
         sensor_positions=sensors,
         B_measured=B_mag,
         B_vectors=B_meas,
         total_volume=total_vol)

print(f"Data saved to {output_filename}")
```
