## Supplementary material for "Modeling the inverse MEG problem in neuro-imaging using Physics Informed Neural Networks": scripts: model_pinns.ipynb

```
# %% [code]
import torch
import torch.nn as nn
import torch.optim as optim
import numpy as np
import os
import matplotlib.pyplot as plt
import matplotlib as mpl
import copy

# Device Configuration
device = torch.device('cuda' if torch.cuda.is_available() else 'cpu')
print(f"Using device: {device}")

# ============================================================================
# SPRINGER PUBLICATION STYLE CONFIGURATION
# ============================================================================
def configure_springer_style():
    """Configure matplotlib for Springer journal submission."""
    mpl.rcParams.update({
        # Fonts
        'font.family': 'sans-serif',
        'font.sans-serif': ['Arial', 'Helvetica', 'DejaVu Sans'],
        'font.size': 9,
        'axes.labelsize': 9,
        'axes.titlesize': 10,
        'xtick.labelsize': 8,
        'ytick.labelsize': 8,
        'legend.fontsize': 8,

        # Lines and Markers
        'lines.linewidth': 1.0,
        'lines.markersize': 3,

        # Figure settings
        'figure.autolayout': True,
        'figure.figsize': (3.3, 2.5), # Default to single column width (3.3 in)
        'figure.dpi': 150,            # Screen PPI

        # Saving
        'savefig.bbox': 'tight',
        'savefig.pad_inches': 0.02,
        'savefig.dpi': 600,           # Publication DPI

        # Colors: Revert to default color cycle (tab10) for color figures
        # 'axes.prop_cycle': plt.cycler(color=['black', 'gray', 'silver', 'black'],
        #                             linestyle=['-', '--', '-', ':'],
        #                             marker=['o', 's', '^', 'd']),
    })

configure_springer_style()

# Helper for sizes
SINGLE_COL_WIDTH = 3.3  # inches (84mm)
DOUBLE_COL_WIDTH = 6.85 # inches (174mm)

def get_figsize(width=SINGLE_COL_WIDTH, aspect=0.75):
    return (width, width * aspect)

# %% [code]
# --- DATA LOADING (Updated for Vector B-Fields & Sensor Positions) ---
# Note: We now assume 'brain_physics_meta.npz' is generated correctly by fenics_1.py
# (i.e. points are already filtered/generated strictly inside the brain).

try:
    from google.colab import files
    if not os.path.exists('meg_training_data.npz'): files.upload()
    if not os.path.exists('brain_physics_meta.npz'): files.upload()
except ImportError:
    pass

data = np.load('meg_training_data.npz')
geo = np.load('brain_physics_meta.npz')

# 1. Load Sensor Positions (From the dataset itself!)
if 'sensors' in data:
    sensor_pos_raw = data['sensors']
    print(f"Loaded Sensor Positions from Dataset: {sensor_pos_raw.shape}")
else:
    # Fallback to file or dummy if strict matching isn't possible (should not happen with new data)
    print("Warning: 'sensors' key not found in .npz. Looking for external file...")
    try:
        sensor_pos_raw = np.load('../sensor_positions.npy')
        print(f"Loaded Sensor Positions from File: {sensor_pos_raw.shape}")
    except FileNotFoundError:
        print("Warning: '../sensor_positions.npy' not found. Using dummy sensors (Mismatches likely!).")
        phi = np.linspace(0, np.pi, 20)
        theta = np.linspace(0, 2*np.pi, 20)
        phi, theta = np.meshgrid(phi, theta)
        x = 0.09 * np.sin(phi) * np.cos(theta)
        y = 0.09 * np.sin(phi) * np.sin(theta)
        z = 0.09 * np.cos(phi)
        sensor_pos_raw = np.stack([x.flatten(), y.flatten(), z.flatten()], axis=1)

# 2. Prepared Training Data
X_raw = data['X']

# Handle Flattening of Vector Fields
if len(X_raw.shape) == 3:
    print(f"Detected Vector B-Fields: {X_raw.shape} -> Flattening to (N, {X_raw.shape[1]*3})")
    X_raw = X_raw.reshape(X_raw.shape[0], -1)
else:
    print(f"Detected Scalar B-Fields: {X_raw.shape}")

# Verify Sensor-Data Alignment
expected_features = sensor_pos_raw.shape[0] * 3
if X_raw.shape[1] != expected_features:
    print(f"CRITICAL WARNING: Feature mismatch! Data has {X_raw.shape[1]} features, but Sensors imply {expected_features}.")
    print("Reconstruction Loss will FAIL. Truncating/Aligning sensors to match data...")
    # Heuristic: If data is 320*3=960, and sensors are 400, cut sensors.
    n_sensors_data = X_raw.shape[1] // 3
    if n_sensors_data < sensor_pos_raw.shape[0]:
        print(f"  -> Truncating sensors from {sensor_pos_raw.shape[0]} to {n_sensors_data}")
        sensor_pos_raw = sensor_pos_raw[:n_sensors_data]

# --- NORMALIZATION STRATEGY (Log-Robust) ---
normalization_type = 'log_robust'

GLOBAL_B_SCALE = 1.0
GLOBAL_B_MEAN = 0.0
GLOBAL_B_STD = 1.0

if normalization_type == 'log_robust':
    # 1. Estimate typical scale
    scale_factor = np.median(np.abs(X_raw))
    if scale_factor == 0: scale_factor = 1e-12

    # 2. Log-Transform
    X_log = np.sign(X_raw) * np.log1p(np.abs(X_raw) / scale_factor)

    # 3. Robust Standard Scaling
    mean_log = np.mean(X_log)
    std_log = np.std(X_log)

    X_norm = (X_log - mean_log) / (std_log + 1e-16)

    # Clip extreme outliers
    X_norm = np.clip(X_norm, -5.0, 5.0)

    # Save stats for later
    GLOBAL_B_SCALE = scale_factor
    GLOBAL_B_MEAN = mean_log
    GLOBAL_B_STD = std_log

    print(f"Input Normalization (Log+Standard): Scale={scale_factor:.2e}, Mean={mean_log:.2f}, Std={std_log:.2f}")

else:
    X_mean, X_std = np.mean(X_raw), np.std(X_raw)
    X_norm = (X_raw - X_mean) / (X_std + 1e-16)
    print(f"Input Normalization (Z-Score): Mean {X_mean:.2e}, Std {X_std:.2e}")

# 3. TARGETS (Geometry): Scale Meters to Normalized Space
Y_raw = data['Y']
max_coords = np.max(np.abs(Y_raw), axis=0) # [x_max, y_max, z_max]
COORD_SCALE_VEC = 1.0 / (max_coords + 1e-8) # Inverse of max extent

print(f"Target Normalization (Vector): Max Extents {max_coords} (m)")
Y_norm = Y_raw * COORD_SCALE_VEC
COORD_SCALE_TENS = torch.tensor(COORD_SCALE_VEC, dtype=torch.float32).to(device)

# Split Train/Val (80/20)
n_total = len(X_norm)
n_train = int(0.8 * n_total)
indices = np.random.RandomState(42).permutation(n_total)

X_train = torch.tensor(X_norm[indices[:n_train]], dtype=torch.float32).to(device)
Y_train = torch.tensor(Y_norm[indices[:n_train]], dtype=torch.float32).to(device)
X_val = torch.tensor(X_norm[indices[n_train:]], dtype=torch.float32).to(device)
Y_val = torch.tensor(Y_norm[indices[n_train:]], dtype=torch.float32).to(device)

# Geometry Points
interior_pts_filtered = geo['interior_pts']
interior_pts_fixed = torch.tensor(interior_pts_filtered, dtype=torch.float32).to(device)
surf_pts = geo['surface_midpoints'] if 'surface_midpoints' in geo else geo['surface_pts']
surf_pts_fixed = torch.tensor(surf_pts, dtype=torch.float32).to(device)
surf_normals_fixed = torch.tensor(geo['surface_normals'], dtype=torch.float32).to(device)

# Sensor Positions to Device
sensor_pos_fixed = torch.tensor(sensor_pos_raw, dtype=torch.float32).to(device)

print(f"Train Set: {len(X_train)} samples | Val Set: {len(X_val)} samples")

# %% [code]
# Calculate the maximum normalized radius from your interior points
with torch.no_grad():
    # norm calculates the distance from (0,0,0) for each point
    brain_radii = torch.norm(interior_pts_fixed * COORD_SCALE_TENS, dim=1)
    MAX_RADIUS = brain_radii.max().item()

print(f"The anatomical boundary (MAX_RADIUS) is: {MAX_RADIUS:.4f}")

# --- UNIFIED MODEL DEFINITION (Shared Encoder) ---
# This architecture uses a 'Y' shape:
# 1. Shared Encoder: Extracts Features (Z) from B-field.
# 2. Location Head: Z -> r (Source Coordinates).
# 3. Potential Head: (Z, x) -> V (Electric Potential).
# This coupling ensures the physics (V) and the solution (r) share a common representation.

class UnifiedMEGPinn(nn.Module):
    def __init__(self, n_sensors, n_hidden=256, latent_dim=128, max_radius=1.8):
        super().__init__()
        self.max_radius = max_radius

        # 1. SHARED ENCODER (B -> Z)
        self.encoder = nn.Sequential(
            nn.Linear(n_sensors, n_hidden), nn.GELU(),
            nn.Linear(n_hidden, n_hidden), nn.GELU(),
            nn.Linear(n_hidden, latent_dim) # Latent Representation
        )

        # 2. LOCATION HEAD (Z -> r)
        self.location_head = nn.Sequential(
            nn.Linear(latent_dim, 64), nn.GELU(),
            nn.Linear(64, 3),
            nn.Tanh() # Squash to [-1, 1]
        )

        # 3. POTENTIAL HEAD (Z + x -> V)
        # Input: Latent (128) + Spatial Coords (3) = 131
        self.potential_head = nn.Sequential(
            nn.Linear(latent_dim + 3, n_hidden), nn.SiLU(), # SiLU for smooth gradients
            nn.Linear(n_hidden, n_hidden), nn.SiLU(),
            nn.Linear(n_hidden, n_hidden), nn.SiLU(),
            nn.Linear(n_hidden, 1)
        )

        # Accessor for optimizer grouping (Legacy adaptation)
        self.inverter = self.location_head # Gradients will flow through encoder too, need careful handling

    def get_latent(self, b_field):
        return self.encoder(b_field)

    def predict_source(self, b_field):
        z = self.encoder(b_field)
        r_norm = self.location_head(z)
        return r_norm * self.max_radius

    def predict_potential(self, inputs):
        # inputs shape: (N_points, latent_dim + 3)
        # Note: This low-level function expects the concatenation to happen outside
        # or we update the API. To keep compatibility with 'compute_physics_loss',
        # we will assume 'inputs' contains [Spatial(3), Latent(128)].
        return self.potential_head(inputs) * 2e-4

# --- API Update ---
# We define the model instantiation here but it will be re-created in training loops.
model = UnifiedMEGPinn(
    n_sensors=X_train.shape[1],
    n_hidden=256,
    latent_dim=128,
    max_radius=MAX_RADIUS
).to(device)


# %% [code]
# --- TRAINING FUNCTION (Updated: Grad Annealing Removed) ---
def train_experiment(name, use_physics=True, epochs=500, lr=1e-3, data_fraction=1.0,
                     n_hidden=256, phy_scale=0.05):
    print(f"\n>>> Starting Experiment: {name}")
    print(f"    Physics Loss: {'ENABLED' if use_physics else 'DISABLED'}")
    if use_physics:
         print(f"    Params: LR={lr:.0e}, Hidden={n_hidden}, PhyScale={phy_scale}")

    # Create Subset
    n_subset = int(len(X_train) * data_fraction)
    print(f"    Training on {n_subset}/{len(X_train)} samples ({data_fraction:.0%} of data)")

    X_train_sub = X_train[:n_subset]
    Y_train_sub = Y_train[:n_subset]

    # Initialize fresh Unified Model
    model = UnifiedMEGPinn(
        n_sensors=X_train.shape[1],
        n_hidden=n_hidden,
        latent_dim=128,          # Using implicit default latent dim
        max_radius=MAX_RADIUS
    ).to(device)

    optimizer = optim.Adam(model.parameters(), lr=lr)
    scheduler = optim.lr_scheduler.ReduceLROnPlateau(optimizer, patience=15, factor=0.5)
    loss_fn = nn.MSELoss()

    history = {
        'train_loss': [], 'val_loss': [], 'val_error_cm': [],
        'loss_data': [], 'loss_phy': [], 'loss_bc': [], 'loss_rec': []
    }

    start_time = time.time()

    for epoch in range(epochs):
        model.train()
        perm = torch.randperm(n_subset)
        epoch_loss = 0

        accum_data = 0.0; accum_phy = 0.0; accum_bc = 0.0; accum_rec = 0.0

        for i in range(0, n_subset, 32):
            idx = perm[i:i+32]
            X_b, Y_b = X_train_sub[idx], Y_train_sub[idx]

            optimizer.zero_grad()
            # Predict r (Forward Pass through B -> Z -> r)
            pred = model.predict_source(X_b)
            loss_data = loss_fn(pred, Y_b)
            accum_data += loss_data.item()

            loss_phy_pure = torch.tensor(0.0).to(device)
            phy_term, bc_term, rec_term = torch.tensor(0.0), torch.tensor(0.0), torch.tensor(0.0)

            if use_physics:
                if epoch >= 10:
                    ramp = min(1.0, (epoch - 10) / 20.0)

                    # Physics Check (Using X_b to recalculate Z internally)
                    # No backward() called inside compute_physics_loss
                    phy_term, bc_term, rec_term = compute_physics_loss(model, pred, X_b)
                    loss_phy_pure = phy_term + bc_term + rec_term

                    # Standard Static Weighting
                    weight = phy_scale * ramp
                    loss = loss_data + weight * loss_phy_pure
                    loss.backward()
                    optimizer.step()

                    epoch_loss += loss.item()
                    accum_phy += (weight * phy_term).item()
                    accum_bc += (weight * bc_term).item()
                    accum_rec += (weight * rec_term).item()
                else:
                    loss_data.backward()
                    optimizer.step()
                    epoch_loss += loss_data.item()
            else:
                loss_data.backward()
                optimizer.step()
                epoch_loss += loss_data.item()

        # Log
        n_batches = max(1, n_subset / 32)
        history['train_loss'].append(epoch_loss / n_batches)
        history['loss_data'].append(accum_data / n_batches)
        history['loss_phy'].append(accum_phy / n_batches)
        history['loss_bc'].append(accum_bc / n_batches)
        history['loss_rec'].append(accum_rec / n_batches)

        # Validation
        model.eval()
        with torch.no_grad():
            pred_val_norm = model.predict_source(X_val)
            val_loss = loss_fn(pred_val_norm, Y_val).item()

            pred_m = pred_val_norm / COORD_SCALE_TENS
            target_m = Y_val / COORD_SCALE_TENS
            diff_m = (pred_m - target_m).cpu().numpy()
            err_cm = np.mean(np.linalg.norm(diff_m, axis=1)) * 100

        scheduler.step(val_loss)
        history['val_loss'].append(val_loss)
        history['val_error_cm'].append(err_cm)

        if epoch % 10 == 0:
            print(f"Ep {epoch:03d} | Data:{history['loss_data'][-1]:.1e} | Rec:{history['loss_rec'][-1]:.1e} | PDE:{history['loss_phy'][-1]:.1e} | Err:{err_cm:.2f}cm")

    duration = time.time() - start_time
    print(f"Done. Duration: {duration:.1f}s | Final Error: {history['val_error_cm'][-1]:.2f} cm")
    return history, model

# %% [code]
def compute_physics_loss(model, pred_source_norm, input_b_norm):
    # Updated for Unified Architecture (Shared Z)

    # 1. Select Random Subset
    batch_size = pred_source_norm.shape[0]
    n_samples = min(batch_size, 4)
    idx = torch.randperm(batch_size)[:n_samples]

    selected_src_norm = pred_source_norm[idx]
    selected_input_b = input_b_norm[idx]

    # NEW: Get Latent Condition for this batch
    # We need 'Z' for the Potential Head
    # Ideally, we should reuse the Z computed during the forward pass to save compute,
    # but for code simplicity here we re-compute or assume access.
    # Since we don't have 'Z' passed in, we re-compute from 'input_b'.
    z_batch = model.get_latent(selected_input_b) # (4, 128)

    # Constants
    SPREAD = 0.005 # Meters
    MOMENT_MAG = 1e-8  # Am
    SIGMA_Cond = 0.33 # S/m
    mu_0 = 4 * np.pi * 1e-7
    NORM_FACTOR = 1.0 / ((SPREAD * np.sqrt(2*np.pi))**3)
    VOL_BRAIN = (4/3) * np.pi * (0.065 * 0.055 * 0.045)

    total_phy, total_bc, total_rec = 0, 0, 0

    xyz_interior = interior_pts_fixed
    xyz_surf = surf_pts_fixed

    for i in range(n_samples):
        src_norm = selected_src_norm[i].unsqueeze(0) # (1, 3)
        src_m = src_norm / COORD_SCALE_TENS        # (1, 3) Meters
        z_sample = z_batch[i].unsqueeze(0)         # (1, 128)

        # --- A. PDE (Poisson) ---
        perm = torch.randperm(xyz_interior.shape[0])[:500]
        xyz_batch_m = xyz_interior[perm].detach().clone()
        xyz_batch_m.requires_grad = True

        # Inputs: [Spatial(3), Latent(128)]
        # Note order must match 'potential_head' definition input layer
        # Layer 0 is Linear(latent + 3 -> ...)
        # We need to stack [Latent, Spatial] or [Spatial, Latent].
        # Looking at cell 3: nn.Linear(latent_dim + 3, ...). usually X comes last or first.
        # Let's standardize on: inputs = cat([z.repeat, x_norm], dim=1) if we assume x is varying

        xyz_batch_norm = xyz_batch_m * COORD_SCALE_TENS

        # Expand Z to match number of spatial points
        z_expanded = z_sample.repeat(len(xyz_batch_m), 1) # (500, 128)

        # Concat: [Latent(128), Spatial(3)] => 131 features
        inputs = torch.cat([z_expanded, xyz_batch_norm], dim=1)

        V = model.predict_potential(inputs)

        # Laplacian (w.r.t Spatial Meters!)
        grad_V = torch.autograd.grad(V.sum(), xyz_batch_m, create_graph=True)[0]
        laplacian = 0
        for j in range(3):
            grad_grad = torch.autograd.grad(grad_V[:, j].sum(), xyz_batch_m, create_graph=True)[0]
            laplacian += grad_grad[:, j]

        # Source Term
        diff = xyz_batch_m - src_m
        dist_sq = torch.sum(diff**2, dim=1).unsqueeze(1)
        gaussian = torch.exp(-torch.clamp(dist_sq, max=0.1)/(2*SPREAD**2)) * NORM_FACTOR
        m_dot_r = MOMENT_MAG * diff[:, 0].unsqueeze(1)
        term_source = (m_dot_r / (SPREAD**2)) * gaussian

        res_pde = (SIGMA_Cond * laplacian.unsqueeze(1)) + term_source
        total_phy += torch.mean(res_pde**2)

        # --- B. Boundary Condition ---
        perm_s = torch.randperm(xyz_surf.shape[0])[:200]
        xyz_s_m = xyz_surf[perm_s].detach().clone()
        xyz_s_m.requires_grad = True
        normals_s = surf_normals_fixed[perm_s]

        # Input Construction
        xyz_s_norm = xyz_s_m * COORD_SCALE_TENS
        z_s_expanded = z_sample.repeat(len(xyz_s_m), 1)
        inputs_s = torch.cat([z_s_expanded, xyz_s_norm], dim=1)

        V_s = model.predict_potential(inputs_s)
        grad_V_s = torch.autograd.grad(V_s.sum(), xyz_s_m, create_graph=True)[0]
        flux = torch.sum(grad_V_s * normals_s, dim=1)
        total_bc += torch.mean(flux**2)

        # --- C. B-Field Reconstruction ---
        # 1. Primary Field (Analytical)
        diff_s = sensor_pos_fixed - src_m
        dist_s = torch.norm(diff_s, dim=1, keepdim=True)
        dist_s_pow3 = torch.clamp(dist_s**3, min=1e-10)

        cross_prod = torch.zeros_like(diff_s)
        cross_prod[:, 1] = -diff_s[:, 2]
        cross_prod[:, 2] =  diff_s[:, 1]
        cross_prod = cross_prod * MOMENT_MAG

        B_prim = (mu_0 / (4*np.pi)) * (cross_prod / dist_s_pow3)

        # 2. Volume Currents (Monte Carlo reuse)
        S = sensor_pos_fixed.unsqueeze(1)
        P = xyz_batch_m.detach().unsqueeze(0)
        diff_vol = S - P
        dist_vol = torch.norm(diff_vol, dim=2, keepdim=True)
        dist_vol_pow3 = torch.clamp(dist_vol**3, min=1e-6)

        G = grad_V.detach().unsqueeze(0)

        cp_x = G[:,:,1]*diff_vol[:,:,2] - G[:,:,2]*diff_vol[:,:,1]
        cp_y = G[:,:,2]*diff_vol[:,:,0] - G[:,:,0]*diff_vol[:,:,2]
        cp_z = G[:,:,0]*diff_vol[:,:,1] - G[:,:,1]*diff_vol[:,:,0]
        CP = torch.stack([cp_x, cp_y, cp_z], dim=2)

        integrand = CP / dist_vol_pow3
        integral = torch.mean(integrand, dim=1) * VOL_BRAIN # (N_sens, 3)

        factor = -(mu_0 * SIGMA_Cond) / (4*np.pi)
        B_vol = factor * integral

        B_total_tesla = B_prim + B_vol
        B_pred_flat = B_total_tesla.flatten()

        # Normalize Pred
        B_pred_log = torch.sign(B_pred_flat) * torch.log1p(torch.abs(B_pred_flat) / GLOBAL_B_SCALE)
        B_pred_norm = (B_pred_log - GLOBAL_B_MEAN) / (GLOBAL_B_STD + 1e-16)

        total_rec += torch.mean((B_pred_norm - selected_input_b[i])**2)

    return total_phy / n_samples, total_bc / n_samples, total_rec / n_samples

# %% [code]
# --- HYPERPARAMETER OPTIMIZATION FOR PINN (Updated: Grad Annealing Removed) ---

def train_pinn_for_hpo(config, epochs=50):
    lr = config.get('lr', 1e-3)
    n_hidden = config.get('n_hidden', 256)
    phy_weight = config.get('phy_weight', 1e-2)

    n_subset = int(len(X_train) * 0.1)
    X_sub = X_train[:n_subset]
    Y_sub = Y_train[:n_subset]

    model = UnifiedMEGPinn(X_train.shape[1], n_hidden=n_hidden, latent_dim=128, max_radius=MAX_RADIUS).to(device)
    optimizer = optim.Adam(model.parameters(), lr=lr)
    loss_fn = nn.MSELoss()

    for epoch in range(epochs):
        model.train()
        perm = torch.randperm(n_subset)

        for i in range(0, n_subset, 32):
            idx = perm[i:i+32]
            X_b, Y_b = X_sub[idx], Y_sub[idx]

            optimizer.zero_grad()
            pred = model.predict_source(X_b)
            loss_data = loss_fn(pred, Y_b)

            performed_step = False

            if epoch >= 5:
                ramp = min(1.0, (epoch - 5) / 10.0)
                phy_term, bc_term, rec_term = compute_physics_loss(model, pred, X_b)
                loss_phy_pure = phy_term + bc_term + rec_term

                # Standard Static Weighting
                loss = loss_data + (phy_weight * ramp) * loss_phy_pure
                loss.backward()
                optimizer.step()
                performed_step = True

            if not performed_step:
                loss_data.backward()
                optimizer.step()

    model.eval()
    with torch.no_grad():
        pred_val_norm = model.predict_source(X_val)
        pred_cm = (pred_val_norm / COORD_SCALE_TENS) * 100.0
        target_cm = (Y_val / COORD_SCALE_TENS) * 100.0
        diff = (pred_cm - target_cm).cpu().numpy()
        err_cm = np.mean(np.linalg.norm(diff, axis=1))

    return err_cm

def run_hyperparameter_optimization(n_trials=10):
    print(f"\n>>> Starting PINN Hyperparameter Search ({n_trials} trials)")
    space = {
        'lr': [1e-3, 5e-4, 2e-3],
        'n_hidden': [256, 512],
        'phy_weight': [0.01, 0.05, 0.1, 0.2]
    }
    results = []
    print(f"{'Trial':<6} | {'LR':<8} | {'Hidden':<8} | {'PhyW':<8} | {'Error (cm)':<10}")
    print("-" * 55)
    for i in range(n_trials):
        config = {
            'lr': np.random.choice(space['lr']),
            'n_hidden': int(np.random.choice(space['n_hidden'])),
            'phy_weight': np.random.choice(space['phy_weight'])
        }
        err = train_pinn_for_hpo(config, epochs=40)
        results.append((err, config))
        print(f"{i+1:<6} | {config['lr']:<8.0e} | {config['n_hidden']:<8} | {config['phy_weight']:<8.0e} | {err:.2f}")

    results.sort(key=lambda x: x[0])
    best = results[0]
    print("\n=== TOP CONFIGURATION ===")
    print(f"Error: {best[0]:.2f} cm")
    print(f"Config: {best[1]}")
    return best[1], results

# 1. Run Optimization
best_config, all_res = run_hyperparameter_optimization(n_trials=12)

# 2. Visualization
errors = [r[0] for r in all_res]
plt.figure(figsize=(10, 6))
plt.barh(range(len(errors)), errors, color='skyblue')
labels = [f"Err: {r[0]:.2f}cm | H:{r[1]['n_hidden']} Lr:{r[1]['lr']:.0e}" for r in all_res]
plt.yticks(range(len(errors)), labels, fontsize=9);
plt.xlabel('Validation Error (cm)'); plt.title('Hyperparameter Random Search Results')
plt.grid(axis='x'); plt.show()

# %% [code]
import time

# EXPERIMENT 1: Full Data (100%)
print("\n=== EXPERIMENT 1: Full Data (100%) ===")
# Using Optimized Parameters found in HPO step
print(f"Using Optimal Config: {best_config}")
hist_pinn_100, final_model = train_experiment("PINN (100%) [Optimized]",
                                    use_physics=True,
                                    epochs=500,
                                    data_fraction=1.0,
                                    n_hidden=best_config['n_hidden'],
                                    lr=best_config['lr'],
                                    phy_scale=best_config['phy_weight'])

# --- LOSS COMPONENT PLOT (Experiment 1) ---
# Springer: Use single column width (3.3 inches) if possible, or double (6.85)
# Loss Components usually need space -> Double column
plt.figure(figsize=(6.85, 4.5), dpi=150)
plt.plot(hist_pinn_100['loss_data'], label='Data Loss (MSE)', linewidth=1)
# Scale Physics components for visibility if they are very small
# Or just plot raw to show their relative magnitude
plt.plot(hist_pinn_100['loss_phy'], label='PDE Loss', linewidth=1, linestyle='--')
plt.plot(hist_pinn_100['loss_bc'], label='Boundary Loss', linewidth=1, linestyle=':')
plt.plot(hist_pinn_100['loss_rec'], label='Reconstruction Loss', linewidth=1, linestyle='-.')
plt.yscale('log')
plt.title('Loss Components (Experiment 1 - Full Data)', fontsize=10)
plt.ylabel('Loss Value (Log Scale)', fontsize=9)
plt.xlabel('Epoch', fontsize=9)
plt.legend(fontsize=8)
plt.grid(True, which="both", ls="-", alpha=0.3)
plt.tight_layout()
plt.show()

# %% [code]


# %% [code]
# Check if best_config exists (from HPO step or Exp 1), if not define robust defaults
if 'best_config' not in locals():
    print("Warning: 'best_config' not found. Using default parameters.")
    best_config = {
        'lr': 1e-3,
        'n_hidden': 256,
        'phy_weight': 0.05
    }

# EXPERIMENT: Scarce Data (10%)
print("\n=== EXPERIMENT : Scarce Data (10%) ===")
# Using Optimized Parameters found in HPO step
print(f"Using Optimal Config: {best_config}")
hist_pinn_10, model_pinn_10 = train_experiment("PINN (10%) [Optimized]",
                                               use_physics=True,
                                               epochs=500,
                                               data_fraction=0.1,
                                               n_hidden=best_config['n_hidden'],
                                               lr=best_config['lr'],
                                               phy_scale=best_config['phy_weight'])


# %% [code]
# --- LOSS COMPONENT PLOT (Experiment on Scarce data) ---
# Use smaller single column for this one if possible, or double
plt.figure(figsize=(3.3, 2.5), dpi=150) # Single Column 84mm
plt.plot(hist_pinn_10['loss_data'], label='Data Loss', linewidth=1)
plt.plot(hist_pinn_10['loss_phy'], label='PDE Loss', linewidth=1, linestyle='--')
plt.plot(hist_pinn_10['loss_bc'], label='Boundary Loss', linewidth=1, linestyle=':')
plt.plot(hist_pinn_10.get('loss_rec', np.zeros_like(hist_pinn_10['loss_data'])), label='Rec Loss', linewidth=1, linestyle='-.')
plt.yscale('log')
plt.title('Loss 10% Data', fontsize=9)
plt.ylabel('Loss (Log Scale)', fontsize=8)
plt.xlabel('Epoch', fontsize=8)
plt.legend(fontsize=7, loc='upper right')
plt.grid(True, which="both", ls="-", alpha=0.3)
plt.tight_layout()
plt.show()


# %% [code]
# --- COMPARATIVE PLOTS ---
# Double column width (174mm = 6.85 in)
plt.figure(figsize=(6.85, 3.0), dpi=150)

# PLOT 1: Full Data (100%)
plt.subplot(1, 2, 1)
if 'hist_pinn_100' in locals():
    plt.plot(hist_pinn_100['val_error_cm'], label='PINN', linewidth=1.5, color='black')
    plt.title('100% Data (Rich)', fontsize=9)
    plt.ylabel('Error (cm)', fontsize=8)
    plt.xlabel('Epoch', fontsize=8)
    plt.legend(fontsize=7)
    plt.grid(True, alpha=0.3)
else:
    plt.text(0.5, 0.5, "Experiment 1 Not Run", ha='center')

# PLOT 2: Scarce Data (10%)
plt.subplot(1, 2, 2)
# Orange/Gray scheme
plt.plot(hist_pinn_10['val_error_cm'], label='PINN (10%)', linewidth=1.5, color='gray', linestyle='--')
plt.title('10% Data (Scarce)', fontsize=9)
plt.ylabel('Error (cm)', fontsize=8)
plt.xlabel('Epoch', fontsize=8)
plt.legend(fontsize=7)
plt.grid(True, alpha=0.3)

plt.tight_layout()
plt.show()

# %% [code]
# --- SEMI-SUPERVISED TRAINING FUNCTION (Simplified: Single Phase) ---
def train_semisupervised(name, labeled_fraction=0.05, epochs=300, lr=1e-3):
    print(f"\n>>> Starting Semi-Supervised Experiment: {name}")

    n_labeled = int(len(X_train) * labeled_fraction)
    X_labeled = X_train[:n_labeled]
    Y_labeled = Y_train[:n_labeled]
    X_unlabeled = X_train[n_labeled:]

    model_pinn = UnifiedMEGPinn(X_train.shape[1], n_hidden=256, latent_dim=128, max_radius=MAX_RADIUS).to(device)

    # Use a scheduler to handle LR dropping automatically
    opt_pinn = optim.Adam(model_pinn.parameters(), lr=lr)
    scheduler = optim.lr_scheduler.ReduceLROnPlateau(opt_pinn, patience=20, factor=0.5)
    loss_fn = nn.MSELoss()
    history = {'pinn_error': []}

    print(f"    Training on {len(X_labeled)} Labeled + {len(X_unlabeled)} Unlabeled samples.")

    for epoch in range(epochs):
        model_pinn.train()

        # 1. PINN on Labeled Data (Supervised)
        perm_l = torch.randperm(len(X_labeled))
        loss_labeled_total = 0

        for i in range(0, len(X_labeled), 32):
            idx = perm_l[i:i+32]
            X_b, Y_b = X_labeled[idx], Y_labeled[idx]
            opt_pinn.zero_grad()
            pred = model_pinn.predict_source(X_b)

            # Physics Check (Using X_b for reconstruction)
            phy_term, bc_term, rec_term = compute_physics_loss(model_pinn, pred, X_b)

            loss = loss_fn(pred, Y_b) + 0.05 * (phy_term + bc_term + rec_term)
            loss.backward()
            opt_pinn.step()
            loss_labeled_total += loss.item()

        # 2. Unlabeled Physics (Unsupervised)
        # We process fewer batches to keep ratio ~1:1 or slightly more unsupervised
        n_unlabeled_batches = min(len(X_unlabeled) // 32, len(X_labeled) // 32 * 2)
        perm_u = torch.randperm(len(X_unlabeled))

        for i in range(n_unlabeled_batches):
            idx = perm_u[i*32 : (i+1)*32]
            X_u = X_unlabeled[idx]
            opt_pinn.zero_grad()
            pred_u = model_pinn.predict_source(X_u)

            # Physics on Unlabeled
            phy_term, bc_term, rec_term = compute_physics_loss(model_pinn, pred_u, X_u)
            loss_phy = phy_term + bc_term + rec_term

            (0.05 * loss_phy).backward()
            opt_pinn.step()

        # Validation
        model_pinn.eval()
        with torch.no_grad():
            pred_pinn = model_pinn.predict_source(X_val)
            val_loss = loss_fn(pred_pinn, Y_val).item()

            diff_pinn = (pred_pinn - Y_val) / COORD_SCALE_TENS
            err_pinn = np.mean(np.linalg.norm(diff_pinn.cpu().numpy(), axis=1)) * 100

        scheduler.step(val_loss)
        history['pinn_error'].append(err_pinn)

        if epoch % 20 == 0:
            print(f"Epoch {epoch:03d} | Labeled Loss: {loss_labeled_total:.1e} | PINN Err: {err_pinn:.2f} cm")

    return history, model_pinn

# RUN EXPERIMENT : Scarce Labels (5%) - Single Long Run
hist_semi, model_semi = train_semisupervised("Semi-Supervised (5% Labels)", labeled_fraction=0.05, epochs=300, lr=1e-3)


# %% [code]
# PLOT RESULTS
# Single column width (3.3 inches)
plt.figure(figsize=(3.3, 2.5), dpi=150)
plt.plot(hist_semi['pinn_error'], label='PINN (5% Labels)', linewidth=1.5)
plt.title('Semi-Supervised Learning', fontsize=9)
plt.ylabel('Error (cm)', fontsize=8)
plt.xlabel('Epoch', fontsize=8)
plt.legend(fontsize=7, loc='upper right')
plt.grid(True, linestyle=":", alpha=0.5)
plt.tight_layout()
plt.show()

# %% [code]
# --- LOSS COMPONENT PLOT (Experiment 1) ---
import matplotlib.pyplot as plt

if 'hist_pinn_100' in locals():
    # Single column
    plt.figure(figsize=(3.3, 2.5), dpi=150)
    plt.plot(hist_pinn_100['loss_data'], label='Data Loss', linewidth=1)
    plt.plot(hist_pinn_100['loss_phy'], label='PDE Loss', linewidth=1, linestyle='--')
    plt.plot(hist_pinn_100['loss_bc'], label='Boundary Loss', linewidth=1, linestyle=':')
    plt.yscale('log')
    plt.title('Loss Components (Full Data)', fontsize=9)
    plt.ylabel('Loss (Log Scale)', fontsize=8)
    plt.xlabel('Epoch', fontsize=8)
    plt.legend(fontsize=7)
    plt.grid(True, which="both", ls="-", alpha=0.3)
    plt.tight_layout()
    plt.show()

    # --- VALIDATION ERROR PLOT ( Experiment 1) ---
    plt.figure(figsize=(3.3, 2.5), dpi=150)
    plt.plot(hist_pinn_100['val_error_cm'], label='Validation Error', linewidth=1.5)
    plt.title('Validation Error (Full Data)', fontsize=9)
    plt.ylabel('Error (cm)', fontsize=8)
    plt.xlabel('Epoch', fontsize=8)
    plt.legend(fontsize=7)
    plt.grid(True, which="both", ls="-", alpha=0.3)
    plt.tight_layout()
    plt.show()

    # --- Summary of Run ---
    completed_epochs = len(hist_pinn_100['val_error_cm'])
    final_val_error = hist_pinn_100['val_error_cm'][-1]
    print(f"\nSummary of  Run:")
    print(f"  Epochs Completed: {completed_epochs}")
    print(f"  Final Validation Error: {final_val_error:.2f} cm")
else:
    print("Error: 'hist_pinn_100' not found. Please ensure Experiment 1 was run  before analyzing.")

# %% [code]
# --- SCATTER COMPARISON (Scarce Data 10% vs Full Data 100%) ---
def plot_scatter_comparison(model, name, ax):
    model.eval()
    with torch.no_grad():
        # Predict (Normalized scale)
        pred_pinn_norm = model.predict_source(X_val).cpu().numpy()
        target_norm = Y_val.cpu().numpy()

        # Unscale to Centimeters (Norm / Scale * 100) -> Norm * (100 / 30)
        # COORD_SCALE_VEC is a vector (3,)
        factor = (1.0 / COORD_SCALE_VEC) * 100.0

        pred_pinn_cm = pred_pinn_norm * factor
        target_cm = target_norm * factor

    # PINN Scatter
    ax.scatter(target_cm[:,0], pred_pinn_cm[:,0], alpha=0.5, label='X', s=20, color='blue', edgecolors='none')
    ax.scatter(target_cm[:,1], pred_pinn_cm[:,1], alpha=0.5, label='Y', s=20, color='orange', edgecolors='none')
    ax.plot([-4, 4], [-4, 4], 'k--', alpha=0.5, linewidth=0.8)
    ax.set_title(name, fontsize=8)
    ax.set_xlabel('True Position (cm)', fontsize=7)
    ax.set_ylabel('Predicted (cm)', fontsize=7)
    ax.tick_params(labelsize=6)
    ax.grid(True, linestyle=":", alpha=0.5)
    ax.legend(fontsize=6, markerscale=2)

# Double column figure (174mm = 6.85in)
plt.figure(figsize=(6.85, 3.0), dpi=150)

# SCATTER 1: Full Data (100%)
ax1 = plt.subplot(1, 2, 1)
if 'final_model' in locals():
    plot_scatter_comparison(final_model, "Accuracy (100% Data)", ax1)
else:
    ax1.text(0.5, 0.5, "Model 1 Not Available", ha='center', fontsize=8)

# SCATTER 2: Scarce Data (10%)
ax2 = plt.subplot(1, 2, 2)
if 'model_pinn_10' in locals():
    plot_scatter_comparison(model_pinn_10, "Accuracy (10% Data)", ax2)
else:
    ax2.text(0.5, 0.5, "Model 10% Not Available", ha='center', fontsize=8)

plt.tight_layout()
plt.show()

# %% [code]
# --- 3D LOCALIZATION VISUALIZATION ---
from mpl_toolkits.mplot3d import Axes3D

def plot_3d_tracking(model, name, X_val, Y_val, ax, n_samples=50):
    model.eval()
    with torch.no_grad():
        pred_full_norm = model.predict_source(X_val).cpu().numpy()
        true_full_norm = Y_val.cpu().numpy()

    # Select random subset
    indices = np.random.choice(len(true_full_norm), n_samples, replace=False)

    # Convert Norm -> CM
    factor = (1.0 / COORD_SCALE_VEC) * 100.0
    pred_sub = pred_full_norm[indices] * factor
    true_sub = true_full_norm[indices] * factor

    # Plot Ground Truth (Green Circles)
    ax.scatter(true_sub[:, 0], true_sub[:, 1], true_sub[:, 2],
               c='blue', marker='o', label='True Source', s=20, alpha=0.6)

    # Plot Predictions (Triangles)

    ax.scatter(pred_sub[:, 0], pred_sub[:, 1], pred_sub[:, 2],
               c='orange', marker='^', label='Pred Source', s=20, alpha=0.8)

    # Connect True -> Pred
    for i in range(len(indices)):  # Use len(indices) instead of n_samples to be safe
        ax.plot([true_sub[i, 0], pred_sub[i, 0]],
                [true_sub[i, 1], pred_sub[i, 1]],
                [true_sub[i, 2], pred_sub[i, 2]],
                color='gray', linestyle=':', alpha=0.5,linewidth=0.5)

    ax.set_title(f"{name}\n(N={n_samples})", fontsize=10)
    ax.set_xlabel('X (cm)', fontsize=8)
    ax.set_ylabel('Y (cm)', fontsize=8)
    ax.set_zlabel('Z (cm)', fontsize=8)
    ax.tick_params(labelsize=7)
    ax.legend(loc='upper right', fontsize=7)

# Create Plot Structure (Double Column)
fig = plt.figure(figsize=(6.85, 5.0), dpi=150)

# PINN
ax1 = fig.add_subplot(111, projection='3d')
plot_3d_tracking(model_pinn_10, "PINN (Scarce Data)", X_val, Y_val, ax1)

plt.suptitle("3D Source Localization", fontsize=11, fontweight='bold')
plt.tight_layout()
plt.show()

# %% [code]
# Select Model to Analyze (Optimized > Default)
if 'final_model' in locals():
    print(">>> Using 'final_model' (Optimized PINN) for analysis.")
    model_to_analyze = final_model
elif 'model_pinn_10' in locals():
    print(">>> Using 'model_pinn_10' for analysis.")
    model_to_analyze = model_pinn_10
else:
    # Fallback for when running just this cell without previous context
    raise NameError("No suitable model found. Please run Training cells first.")

model_to_analyze.eval()

with torch.no_grad():
    # 3. Get the predicted source locations for X_val
    pred_pinn_norm = model_to_analyze.predict_source(X_val).cpu().numpy()

    # 4. Convert normalized predicted and true source locations to centimeters
    # The conversion factor is (1.0 / COORD_SCALE_VEC) * 100.0 from numpy array
    factor = (1.0 / COORD_SCALE_VEC) * 100.0
    pred_pinn_cm = pred_pinn_norm * factor
    target_cm = (Y_val.cpu().numpy()) * factor

    # 5. Calculate the difference between pred_pinn_cm and target_cm
    diff_cm = pred_pinn_cm - target_cm

    # 6. Compute the Euclidean distance (L2 norm) for each difference vector
    individual_errors_cm = np.linalg.norm(diff_cm, axis=1)

print(f"Calculated individual errors for {len(individual_errors_cm)} validation points.")
print(f"First 5 individual errors (cm): {individual_errors_cm[:5]}")
print(f"Mean individual error (cm): {np.mean(individual_errors_cm):.2f}")

# %% [markdown]
**Reasoning**:
Now that the individual errors have been calculated, the next step is to identify the 10 points with the largest errors to prepare for visualization. This involves sorting the errors and selecting the corresponding indices from the validation set.


# %% [code]
num_top_errors = 10

# Sort individual_errors_cm in descending order and get the indices
sorted_indices = np.argsort(individual_errors_cm)[::-1]

# Get the indices of the 10 largest errors
top_10_error_indices = sorted_indices[:num_top_errors]

# Retrieve the true and predicted locations for these top 10 error points
true_top_10_cm = target_cm[top_10_error_indices]
pred_top_10_cm = pred_pinn_cm[top_10_error_indices]
top_10_errors_values = individual_errors_cm[top_10_error_indices]

print(f"Identified {num_top_errors} points with the largest localization errors.")
print("Top 10 errors (cm) and their corresponding true/predicted locations:")
for i in range(num_top_errors):
    print(f"  Error: {top_10_errors_values[i]:.2f} cm | True: {true_top_10_cm[i]} | Pred: {pred_top_10_cm[i]}")

# %% [code]
# --- SAVE MODEL (Compatible with Colab & Local) ---
import os
import torch

model_filename = 'pinn_meg_experiment_1_final.pth'

# 1. Save to the current environment (Local disk or Colab VM)
if 'final_model' in locals():
    torch.save(final_model.state_dict(), model_filename)
    print(f"✅ Model saved to environment: {os.path.abspath(model_filename)}")
else:
    print("⚠️ Warning: 'final_model' variable not found. Did Experiment 1 finish?")

# 2. If running in Google Colab, trigger a browser download
try:
    from google.colab import files
    print("\n[Google Colab Detected]")
    print(f"⬇️ Triggering download for {model_filename}...")
    files.download(model_filename)
except ImportError:
    print("\n[Local Execution Detected]")
    print(f"   Model is saved in your current folder: {os.getcwd()}")

# 3. Optional: Save to Google Drive (Uncomment if needed in Colab)
# from google.colab import drive
# drive.mount('/content/drive')
# shutil.copy(model_filename, '/content/drive/My Drive/pinn_meg_experiment_1_final.pth')
```
