## Supplementary material for "Modeling the inverse MEG problem in neuro-imaging using Physics Informed Neural Networks": scripts: solve_with_mne_inverse.py

```
import numpy as np
import meshio
import time
import os
import matplotlib.pyplot as plt

def run_optimized_mne(data_file='fenics_poisson_forward_data.npz', 
                      stl_file='MalinBjornsdotterBrain55mm.stl',
                      n_sources=20000, 
                      snr=100.0):
    
    print("=================================================================")
    print("   OPTIMIZED MEG INVERSE PROBLEM (MNE + ANATOMICAL CONSTRAINTS)")
    print("=================================================================")

    # 1. LOAD DATA
    use_training_data_format = False
    
    if 'meg_training_data.npz' in data_file: 
        print(f"  > Detected Training Data Format: {data_file}")
        use_training_data_format = True
        
    if not os.path.exists(data_file):
        print(f"Error: {data_file} not found.")
        return

    data = np.load(data_file)
    
    if use_training_data_format:
        # Load logic for meg_training_data.npz (X, Y, sensors)
        # Select one sample to invert (e.g., the last one or a random one)
        # We pick index -1 to be consistent with 'latest'
        idx = -1 
        
        sensor_positions = data['sensors']
        B_vectors_all = data['X'] # (N, 320, 3)
        dipole_pos_all = data['Y'] # (N, 3)
        
        B_vectors = B_vectors_all[idx] # (320, 3)
        dipole_pos_true = dipole_pos_all[idx]
        
        B_meas_clean = B_vectors.flatten()
        use_vector_data = True
        
        # Radii info might be missing in training data, defaulting to standard
        radii_m = np.array([0.065, 0.055, 0.045]) 
        center_m = np.zeros(3) # Training data uses centered mesh
        
        print(f"  > Selected Sample Index: {len(B_vectors_all) + idx}")
    else:
        # Legacy/Original logic for fenics_poisson_forward_data.npz
        sensor_positions = data['sensor_positions']
        
        # CRITICAL: Use Vector Field Data if available
        # Magnitude-only inverse is ambiguous and non-linear.
        # Linear Inverse requires component data.
        if 'B_vectors' in data:
            print("  > Using Vector Field Data (Bx, By, Bz)")
            B_vectors = data['B_vectors'] # (n_sens, 3)
            # Flatten: [Bx1, By1, Bz1, Bx2, By2, Bz2, ...]
            B_meas_clean = B_vectors.flatten() # (n_sens * 3,)
            use_vector_data = True
        else:
            print("  > Warning: Only Magnitude data available. Using scalar approx.")
            B_meas_clean = data['B_measured']
            use_vector_data = False
            
        dipole_pos_true = data['dipole_pos']
        radii_m = data['radii_m']
        center_m = np.zeros(3)
    
    # Add noise
    np.random.seed(42)
    noise_std = np.max(np.abs(B_meas_clean)) / snr
    noise = np.random.randn(len(B_meas_clean)) * 0.01 * np.max(np.abs(B_meas_clean))
    B_meas = B_meas_clean + noise
    
    print(f"Data Loaded: {len(sensor_positions)} sensors")
    if use_vector_data:
        print(f"Measurements: {len(B_meas)} (Vector components)")
    else:
        print(f"Measurements: {len(B_meas)} (Scalar magnitudes)")
        
    print(f"True Dipole: {dipole_pos_true * 100} cm")
    
    # 2. DEFINE SOURCE SPACE (MULTI-LAYER VOLUMETRIC - MATCHES BENCHMARK)
    start_time = time.time()
    source_space = None
    
    if os.path.exists(stl_file):
        print(f"Loading Anatomical Mesh: {stl_file}...")
        mesh = meshio.read(stl_file)
        # Convert mm to meters and center
        # CRITICAL: Must match FEniCS forward model centering exactly
        raw_pts = mesh.points * 1e-3
        
        # FEniCS script used: center = points.mean(axis=0)
        center_offset = np.mean(raw_pts, axis=0)
        
        mesh_verts = raw_pts - center_offset
        
        # MULTI-LAYER VOLUMETRIC SOURCE SPACE (EXACT BENCHMARK METHOD)
        # 5 layers from surface to deep, 4000 sources per layer = 20k total
        print(f"  > Generating multi-layer volumetric source space...")
        layers = [1.0, 0.9, 0.8, 0.7, 0.6]
        n_per_layer = n_sources // len(layers)
        source_layers = []
        
        for scale in layers:
            idx = np.random.choice(len(mesh_verts), n_per_layer, replace=True)
            layer_pts = mesh_verts[idx] * scale  # Scale inward
            source_layers.append(layer_pts)
        
        source_space = np.vstack(source_layers)
    else:
        print("Warning: STL not found. Using volumetric ellipsoidal grid...")
        # Fallback to ellipsoidal volumetric grid (benchmark method)
        source_space = []
        while len(source_space) < n_sources:
            p = (np.random.rand(3) * 2 - 1) * radii_m * 0.9 + center_m
            if np.sum(((p - center_m) / radii_m)**2) <= 0.9:
                source_space.append(p)
        source_space = np.array(source_space)

    print(f"  > Source Space: {len(source_space)} dipoles (Multi-layer volumetric)")
    
    # Note: We don't snap to mesh since volumetric approach should naturally include the dipole location

    # 3. COMPUTE LEAD FIELD (Fully Vectorized)
    # G shape depends on data type
    print("Computing Lead Field Matrix (Biot-Savart)...")
    n_sens = len(sensor_positions)
    n_src = len(source_space)
    
    # Coordinates arrays
    # r_sens: (n_sens, 1, 3)
    # r_src:  (1, n_src, 3)
    r_sens = sensor_positions[:, np.newaxis, :]
    r_src = source_space[np.newaxis, :, :]
    
    # Vector from src to sens: R = r_sens - r_src
    # Shape: (n_sens, n_src, 3)
    R = r_sens - r_src
    dist = np.linalg.norm(R, axis=2) # (n_sens, n_src)
    
    # Constants
    mu0 = 4 * np.pi * 1e-7
    factor = mu0 / (4 * np.pi)
    
    inv_dist3 = 1.0 / (dist**3) # (n_sens, n_src)
    
    if use_vector_data:
        # G shape: (n_sens * 3, n_src * 3)
        # We construct rows for Bx, By, Bz
        G = np.zeros((n_sens * 3, n_src * 3))
        
        # Orientations Qx=[1,0,0], Qy=[0,1,0], Qz=[0,0,1]
        
        # For Qx source (col stride 0):
        # B = [0, -Rz, Ry] * factor / d^3
        # Rows:
        # Bx (row stride 0): 0
        # By (row stride 1): -Rz
        # Bz (row stride 2): Ry
        
        # Row indices
        rows_x = np.arange(0, n_sens*3, 3)
        rows_y = np.arange(1, n_sens*3, 3)
        rows_z = np.arange(2, n_sens*3, 3)
        
        # --- SRC X (cols 0, 3, 6...) ---
        G[rows_x, 0::3] = 0.0
        G[rows_y, 0::3] = -R[:,:,2] * factor * inv_dist3
        G[rows_z, 0::3] =  R[:,:,1] * factor * inv_dist3
        
        # --- SRC Y (cols 1, 4, 7...) ---
        # Q=[0,1,0] -> B = [Rz, 0, -Rx]
        G[rows_x, 1::3] =  R[:,:,2] * factor * inv_dist3
        G[rows_y, 1::3] = 0.0
        G[rows_z, 1::3] = -R[:,:,0] * factor * inv_dist3
        
        # --- SRC Z (cols 2, 5, 8...) ---
        # Q=[0,0,1] -> B = [-Ry, Rx, 0]
        G[rows_x, 2::3] = -R[:,:,1] * factor * inv_dist3
        G[rows_y, 2::3] =  R[:,:,0] * factor * inv_dist3
        G[rows_z, 2::3] = 0.0
        
    else:
        # Scalar Magnitude approx
        # We need G for 3 orientations (x, y, z) per source
        G = np.zeros((n_sens, n_src * 3))
        
        # X-dipoles (Q = [1, 0, 0]) -> cross(Q, R) = [0, -Rz, Ry]
        Bx_x = np.zeros((n_sens, n_src))
        Bx_y = -R[:, :, 2]
        Bx_z =  R[:, :, 1]
        B_x_vec = np.stack([Bx_x, Bx_y, Bx_z], axis=2) * factor * inv_dist3[:, :, np.newaxis]
        G[:, 0::3] = np.linalg.norm(B_x_vec, axis=2)
        
        # Y-dipoles (Q = [0, 1, 0]) -> cross(Q, R) = [Rz, 0, -Rx]
        By_x =  R[:, :, 2]
        By_y =  np.zeros((n_sens, n_src))
        By_z = -R[:, :, 0]
        B_y_vec = np.stack([By_x, By_y, By_z], axis=2) * factor * inv_dist3[:, :, np.newaxis]
        G[:, 1::3] = np.linalg.norm(B_y_vec, axis=2)
        
        # Z-dipoles (Q = [0, 0, 1]) -> cross(Q, R) = [-Ry, Rx, 0]
        Bz_x = -R[:, :, 1]
        Bz_y =  R[:, :, 0]
        Bz_z =  np.zeros((n_sens, n_src))
        B_z_vec = np.stack([Bz_x, Bz_y, Bz_z], axis=2) * factor * inv_dist3[:, :, np.newaxis]
        G[:, 2::3] = np.linalg.norm(B_z_vec, axis=2)
    
    print("  > Lead Field Computed.")

    # --- ENHANCEMENT: DEPTH WEIGHTING ---
    # Standard MNE bias is towards superficial sources.
    # We re-weight G columns by ||g_i||^-p (p=0.8 usually)
    print("Applying Depth Weighting...")
    # Norm of each column (source gain)
    col_norms = np.linalg.norm(G, axis=0)
    # Avoid zero division
    col_norms = np.clip(col_norms, 1e-12, None)
    
    # For vector sources, we often weight the triplet (x,y,z) by the same factor
    # Average norm over the triplet
    triplet_norms = np.zeros(n_src)
    for i in range(n_src):
        # Taking max sensitivity of the triplet
        triplet_norms[i] = np.max(col_norms[i*3:i*3+3])
        
    depth_limit = 0.8 # Standard MNE value
    
    R_diag = np.zeros(n_src * 3)
    for i in range(n_src):
        # Weight for this location
        w_loc = (1.0 / triplet_norms[i]) ** depth_limit 
        R_diag[i*3:i*3+3] = w_loc
        
    # 4. INVERSE SOLUTION (Weighted MNE - EXACT BENCHMARK METHOD)
    print(f"  > Solving Inverse (SNR={snr}, lambda2={1.0/snr**2:.2e})...")
    
    if use_vector_data:
        n_measurements = n_sens * 3
    else:
        n_measurements = n_sens
        
    # Regularization: EXACT BENCHMARK VALUE
    SNR = snr
    lambda2 = 1.0 / SNR**2
    
    # Data Covariance: G R G^T + lambda C_noise
    # R is diagonal, so G R G^T is weighted sum of outer products of columns
    # Efficient: (G * sqrt(R)) @ (G * sqrt(R)).T
    
    sqrt_R_diag = np.sqrt(R_diag)
    G_weighted = G * sqrt_R_diag[np.newaxis, :] # Broadcasting scale columns
    
    Gram = G_weighted @ G_weighted.T
    Cov = Gram + lambda2 * np.eye(n_measurements)
    
    # Solve Cov * w = B (EXACT BENCHMARK METHOD)
    try:
        w = np.linalg.solve(Cov, B_meas)
    except np.linalg.LinAlgError:
        w = np.linalg.lstsq(Cov, B_meas, rcond=1e-5)[0]
    
    # J = R G.T w
    J_est_weighted = G_weighted.T @ w
    J_est = J_est_weighted * sqrt_R_diag # Scale back to physical currents
    
    # 5. LOCALIZE PEAK (EXACT BENCHMARK METHOD)
    J_vectors = J_est.reshape(n_src, 3)
    J_magnitudes = np.linalg.norm(J_vectors, axis=1)
    
    peak_idx = np.argmax(J_magnitudes)
    peak_pos = source_space[peak_idx]
    peak_moment = J_vectors[peak_idx]
    
    error = np.linalg.norm(peak_pos - dipole_pos_true)
    
    elapsed = time.time() - start_time
    
    print("\n-----------------------------------------------------------------")
    print(f"RESULTS:")
    print(f"  True Pos:   {dipole_pos_true * 100} cm")
    print(f"  Est Pos:    {peak_pos * 100} cm")
    print(f"  Error:      {error * 100:.4f} cm")
    print(f"  Time:       {elapsed:.2f} s")
    print("\n-----------------------------------------------------------------")
    
    # Visualize top 1% sources if error is low
    threshold = np.percentile(J_magnitudes, 99.5)
    active_idcs = J_magnitudes > threshold
    active_pts = source_space[active_idcs]
    
    if error * 100 < 1.0:
        print("SUCCESS: Error is under 1.0 cm!")
    else:
        print("Note: Error above 1.0 cm may be expected for volumetric source space.")

    return peak_pos, peak_moment, J_vectors, active_pts, elapsed

if __name__ == "__main__":
    # Run with EXACT BENCHMARK PARAMETERS
    # Multi-layer volumetric: 20k sources across 5 depth layers
    # High SNR for synthetic data: SNR=100.0
    
    # Check if training data exists to ensure consistency with FEniCS/PINN scripts
    if os.path.exists('meg_training_data.npz'):
        print("Using meg_training_data.npz for consistency with FEniCS 2 & PINNs v25...")
        run_optimized_mne(data_file='meg_training_data.npz', n_sources=20000, snr=100.0)
    else:
        print("Warning: meg_training_data.npz not found. Using default fenics_poisson_forward_data.npz")
        run_optimized_mne(n_sources=20000, snr=100.0)
```
